# RNA base pairing complexity in living cells visualized by correlated chemical probing

**DOI:** 10.1101/596353

**Authors:** Anthony M. Mustoe, Nicole Lama, Patrick S. Irving, Samuel W. Olson, Kevin M. Weeks

## Abstract

RNA structure and dynamics are critical to biological function. However, strategies for determining RNA structure *in vivo* are limited, with established chemical probing and newer duplex detection methods each having notable deficiencies. Here we convert the common reagent dimethyl sulfate (DMS) into a useful probe of all four RNA nucleotides. Building on this advance, we introduce PAIR-MaP, which uses single-molecule correlated chemical probing to directly detect base pairing interactions in cells. PAIR-MaP has superior resolution and accuracy compared to alternative experiments, can resolve alternative pairing interactions of structurally dynamic RNAs, and enables highly accurate structure modeling, including of RNAs containing multiple pseudoknots and extensively bound by proteins. Application of PAIR-MaP to human RNase MRP and two bacterial mRNA 5'-UTRs reveals new functionally important and complex structures undetectable by conventional analyses. PAIR-MaP is a powerful, experimentally concise, and broadly applicable strategy for directly visualizing RNA base pairs and dynamics in cells.

## INTRODUCTION

RNA molecules are strongly driven to fold back on themselves into base-paired secondary structures. These structures play central roles in RNA biology, from mediating complex functions such as RNA catalysis and specific ligand recognition, to more broadly tuning RNA sequence accessibility to regulate processes such as translation initiation (1, 2). Furthermore, many RNAs fold into multiple structures, providing the basis for molecular switching functions (3). Accurately resolving RNA structure and its potential dynamic complexity is therefore essential for understanding RNA function.

Chemical probing experiments are among the most broadly useful classes of experiments for characterizing RNA structure (4-6). SHAPE reagents, dimethyl sulfate (DMS), or other chemical probes are used to selectively modify conformationally flexible nucleotides and reactivity is measured using sequencing approaches such as mutational profiling (MaP) (7). These reactivity data provide powerful insight into local RNA structure and can be used to guide accurate RNA structure modeling (7-9). Nevertheless, chemical probing experiments are limited in that they do not directly detect RNA base pairing interactions – structure can only be *inferred* based on compatibility with reactivity data. In some cases, the reactivity data may be equally compatible with multiple structures. Even if the structure inference problem is uniquely defined, follow-up mutational analysis is often desired to obtain *direct* evidence of pairing interactions. Chemical probing data are also poorly suited for resolving alternative structural states of dynamic RNAs. Finally, conventional chemical probing data are difficult to interpret for RNAs bound by proteins or in cells.

To address the limitations of chemical probing experiments, new strategies have been developed that use scanning mutagenesis and chemical probing (mutate-and-map) to identify interacting nucleotides (10) or detect RNA duplexes by crosslinking and proximity ligation (11, 12). However, both of these classes of experiments are laborious, with the former limited to *in vitro* settings and the latter having poorly benchmarked accuracy, low resolution (10-20 nts), and insufficient information to rank and define complete RNA structures (13). We recently introduced a third strategy that uses single-molecule chemical probing experiments (14) to detect correlated modifications between paired nucleotides (15), but this approach was also limited to *in vitro* settings and the underlying mechanism has been questioned (10). Thus, current duplex detection strategies retain substantial limitations, being restricted either to *in vitro* contexts or lacking the desired quantitative accuracy and experimental concision.

Here, we introduce a new strategy that converts the classic reagent DMS into a reliable probe of all four RNA nucleotides. We combine this advance with new analysis algorithms to demonstrate that single-molecule correlated chemical probing reliably detects RNA duplexes in cells, comprising a strategy we term PAIR-MaP (<u>P</u>airing <u>a</u>scertained from interacting <u>R</u>NA strands measured by <u>m</u>utational <u>p</u>rofiling). PAIR-MaP permits simultaneous measurement of local chemical probing data and duplex interactions via one straightforward chemical probing experiment, enabling highly accurate RNA structure modeling and revealing alternative RNA structural states. Application of PAIR-MaP to human RNase MRP and the *E. coli* S2- and S4-binding autoregulatory elements reveals new functionally important structural features of these RNAs, highlighting the broad potential of PAIR-MaP for understanding RNA biology.

## RESULTS

### DMS reliably probes structure of all four nucleotides

DMS is among the most commonly used RNA chemical probes, favored for its cell-permeability and ability to heavily modify RNA molecules during correlated chemical probing experiments. However, a major limitation is that DMS does not typically react with the base pairing face of guanosine (G) and uridine (U) nucleotides due to protonation of the respective N1 and N3 positions at neutral pH (pKa ≈ 9.2; Fig. 1A) (4, 16). We discovered that DMS can be converted into a useful probe of all four nucleotides by performing modification at pH 8, which promotes transient deprotonation of G and U and reaction with DMS. Optimized buffer conditions consisting of 200 mM bicine at pH 8.0 were found to maintain a well-controlled pH without quenching the DMS reaction (SI Methods). These optimized conditions were used to perform multiple-hit DMS probing of natively extracted (termed cell-free) total *E. coli* RNA, and DMS methylation sites were detected using the single-molecule MaP strategy (14). Analysis of the 16S and 23S ribosomal RNAs (rRNAs) reveals that U and G nucleotides are consistently modified in a structure-specific manner: single-stranded U and G positions are modified at average rates of 1.3% and 0.7%, respectively, whereas paired positions are protected and have approximately 4-fold lower modification rates (Fig. 1B, C). The modification rate for U and G residues is ∼10-fold lower than that for A and C (Fig. 1B), but definitively exceeds the threshold required for reliable quantification by the MaP strategy (14).

**Figure 1:**
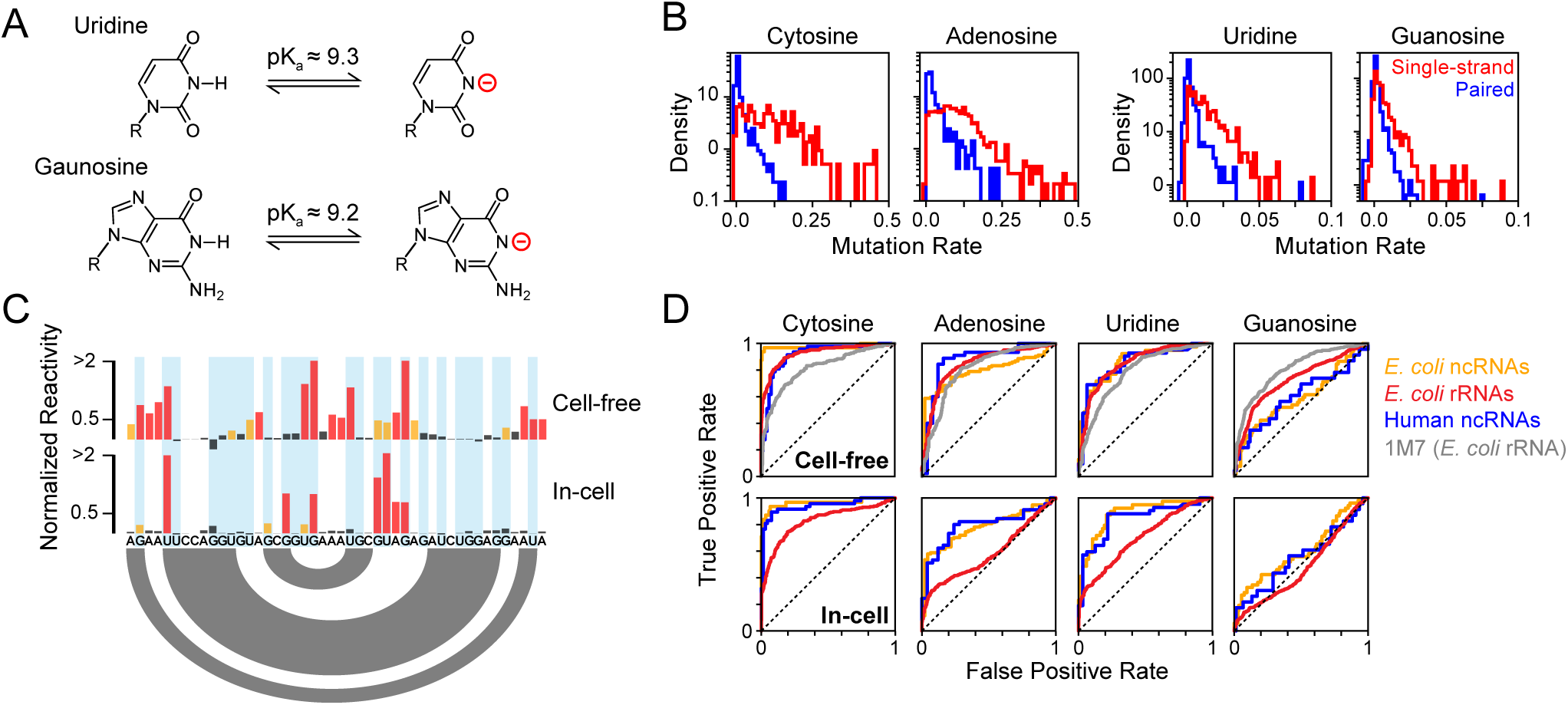
DMS probes all four RNA nucleotides. (**A**) Deprotonation equilibrium of G and U nucleotides (16). (**B**) DMS modification rates measured by MaP for *E. coli* 16S and 23S rRNAs probed under cell-free conditions in buffered bicine (pH 8.0). (**C**) Normalized DMS reactivities for a representative region of the 16S rRNA (nucleotides 693 to 718). The secondary structure is indicated by arcs (bottom). U and G nucleotides are highlighted (blue). (**D**) Receiver Operating Characteristic (ROC) curves for DMS-MaP reactivity profiles calculated for different RNAs. Area under the ROC curve (AUC) values are provided in Table S1. *E. coli* rRNAs: 5S, 16S, and 23S rRNAs. *E. coli* ncRNAs: RNase P and tmRNA. Human ncRNAs: U1 snRNA and RMRP. Cell-free 1M7 SHAPE data from the *E. coli* 16S and 23S rRNA is provided as a reference (17).

Benchmarking across a diverse panel of RNAs with known structures confirmed that DMS reactivity at U and G residues provides a reliable measurement of nucleotide pairing status. In addition to the 16S and 23S rRNA, we used MaP to quantify DMS modification of 5S rRNA, RNase P, and tmRNA in cell-free *E. coli* RNA. We also performed DMS probing experiments on cell-free total RNA from human Jurkat cells and quantified modifications of U1 snRNP and RNase MRP (RMRP). Remarkably, DMS reactivity discriminates single-stranded versus paired U residues with accuracy comparable as for A and C nucleotides and also performs comparably to SHAPE reactivity (Fig. 1D). DMS reactivity is less discriminative for G nucleotides but is still informative (Fig. 1D; Table S1). The decreased specificity observed for G modifications is most likely attributable to non-specific DMS modification at the N7 position of G (4) that is partially detected by MaP.

We also assessed whether DMS is an effective probe of G and U nucleotides in cells. Living *E. coli* or human Jurkat cultures were supplemented with bicine probing buffer and treated with DMS. As expected, DMS is less effective at discriminating single-stranded versus paired nucleotides in cells due to protection by proteins, particularly for the *E. coli* rRNAs (Fig. 1C, D). Nonetheless, DMS still measures structure-specific modification of U nucleotides in cells in all RNAs, again with similar discriminatory power as for A and C nucleotides (Fig. 1C, 1D). DMS reactivity at G nucleotides is weakly informative for *E. coli* and human non-coding RNA structure but is uninformative in the highly protected *E. coli* rRNA.

Combined, our data clearly show that DMS is an effective probe of all four RNA nucleotides at pH 8.0, including in living bacterial and human cells. Separately, our data also demonstrate that the MaP strategy, in conjunction with the *ShapeMapper* bioinformatics pipeline (17), detects DMS modifications with excellent structural specificity without need for specialized enzymes or separate counting of termination events (18, 19).

### PAIR-MaP enables direct visualization of RNA base pairing complexity

The ability to probe all four nucleotides with DMS is an important experimental innovation but does not address the core limitation of conventional RNA structure probing analysis: structures are not visualized directly, but only inferred based on consistency with a one-dimensional reactivity profile. A unique advantage of MaP compared to alternative “seq” readout strategies is that it allows measurement of multiple, correlated DMS modifications within a single RNA molecule (14). We previously showed that we could use correlated chemical probing to detect correlated modifications that occur between A-U and G-C base pairs in model *in vitro* transcripts (15). However, we were unable to detect base pairs in endogenous RNAs due to low DMS reactivity at G and U positions. We now exploit PAIR-MaP to directly detect pairing interactions in endogenous RNAs, including in living cells, at high resolution and excellent specificity.

PAIR-MaP is predicated on detecting correlated DMS modifications on opposing strands of paired duplexes (Fig. 2A). While paired nucleotides are normally protected, equilibrium fluctuations transiently expose paired bases, mediating low but detectable rates of DMS modification. Chance modification of one base will permanently destabilize the base pair, increasing the probability of subsequent DMS modification at either the directly opposing base or neighboring bases (Fig. 2A). We detect these characteristic correlated modification signals by performing correlation analysis over 3-nt windows, which amplifies the weak modification signals of paired nucleotides by summing over nearest-neighbors. Paired duplexes can then be specifically identified as lowly reactive, complementary 3-nt windows that are modified in a correlated manner (Fig. 2B, S1). Significantly, PAIR-MaP detects duplexes formed in the predominant structure of an RNA as well as duplexes formed in lesser but appreciably populated alternative or misfolded structures. We therefore classify PAIR-MaP correlations into two classes (Fig. 2B, S1). “Principal” correlations are defined to occur between lowly reactive positions and are unambiguously the strongest correlation for each set of interacting nucleotides, providing high-confidence indicators of the predominant structure. “Minor” correlations represent weaker correlations or occur between moderately reactive nucleotides and report on unstable and alternative RNA duplexes.

**Figure 2:**
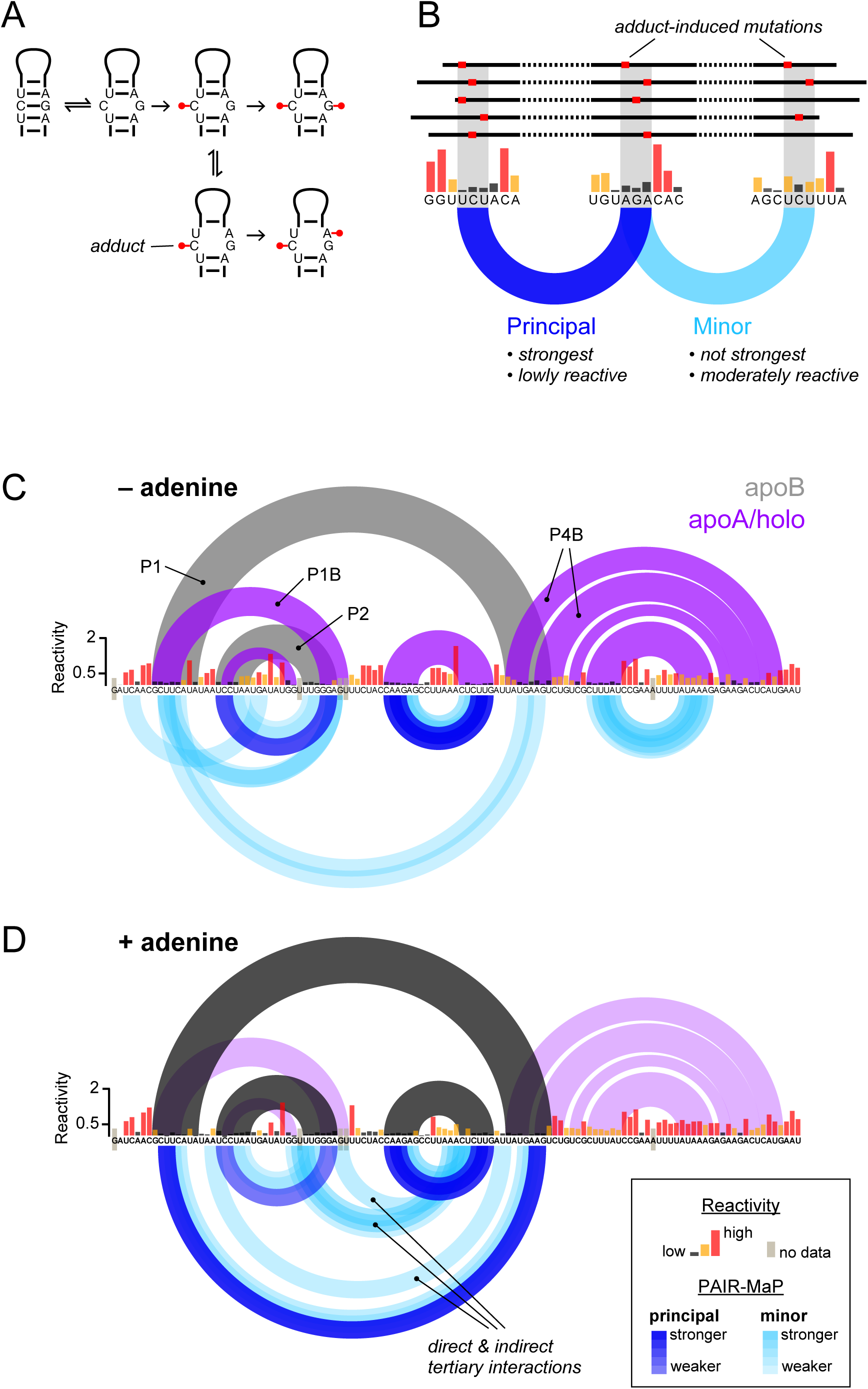
PAIR-MaP enables direct detection of principal and alternative base pairing interactions. (**A**) Correlated modification mechanism. (**B**) Strategy for specifically identifying base pairing interactions from correlated probing data. (**C, D**) *In vitro* PAIR-MaP data collected on the adenine riboswitch (C) without adenine and (D) in the presence of 100 μM adenine. The known secondary structures are shown at top and PAIR-MaP data are shown at bottom.

As an initial validation of our strategy, we used PAIR-MaP to probe an *in vitro* transcript of the *V. vulnificus add* adenine riboswitch, an established model system known to adopt multiple structures (Fig. 2C) (20, 21). PAIR-MaP immediately reveals the complex structural landscape of the riboswitch. In the absence of adenine ligand, PAIR-MaP reports a superposition of multiple pairing interactions recapitulating the ligand-free aptamer (*apoA*) and alternative structure (*apoB*) equilibrium (Fig. 2C). The relative strengths of the P1, P1B, and P2 helix correlations are also consistent with reported stabilities of these helices (populations between 20%-50%) (20, 21). Upon addition of the adenine ligand, the PAIR-MaP correlation network markedly consolidates. All *apoB*-specific correlations disappear, consistent with the expected depopulation of the *apoB* state (expected population <20%), while P1 correlations significantly strengthen (expected population ∼80%) (20, 21). We also observe several minor correlations arising from tertiary interactions and indirect cooperative folding interactions, representing false positive base pairs (but true tertiary interactions). Thus, other types of structural correlations can occasionally pass through the PAIR-MaP filtering algorithm. Combined, these data validate PAIR-MaP as a sensitive and specific strategy for directly visualizing RNA base pairing and structural complexity.

### Direct visualization of RNA base pairing in cells

We next benchmarked PAIR-MaP using endogenous *E. coli* and human RNAs, probed in the cell-free state. PAIR-MaP again provides a detailed visualization of the architectures of these diverse RNAs. For the 16S and 23S rRNAs, extensive correlations clearly define individual domains, including numerous duplexes spanning >350 nucleotides (Fig. 3, S2-S4). PAIR-MaP correlations also clearly define pseudoknots in tmRNA and RNase P (Fig. 3C, S2, S3).

**Figure 3:**
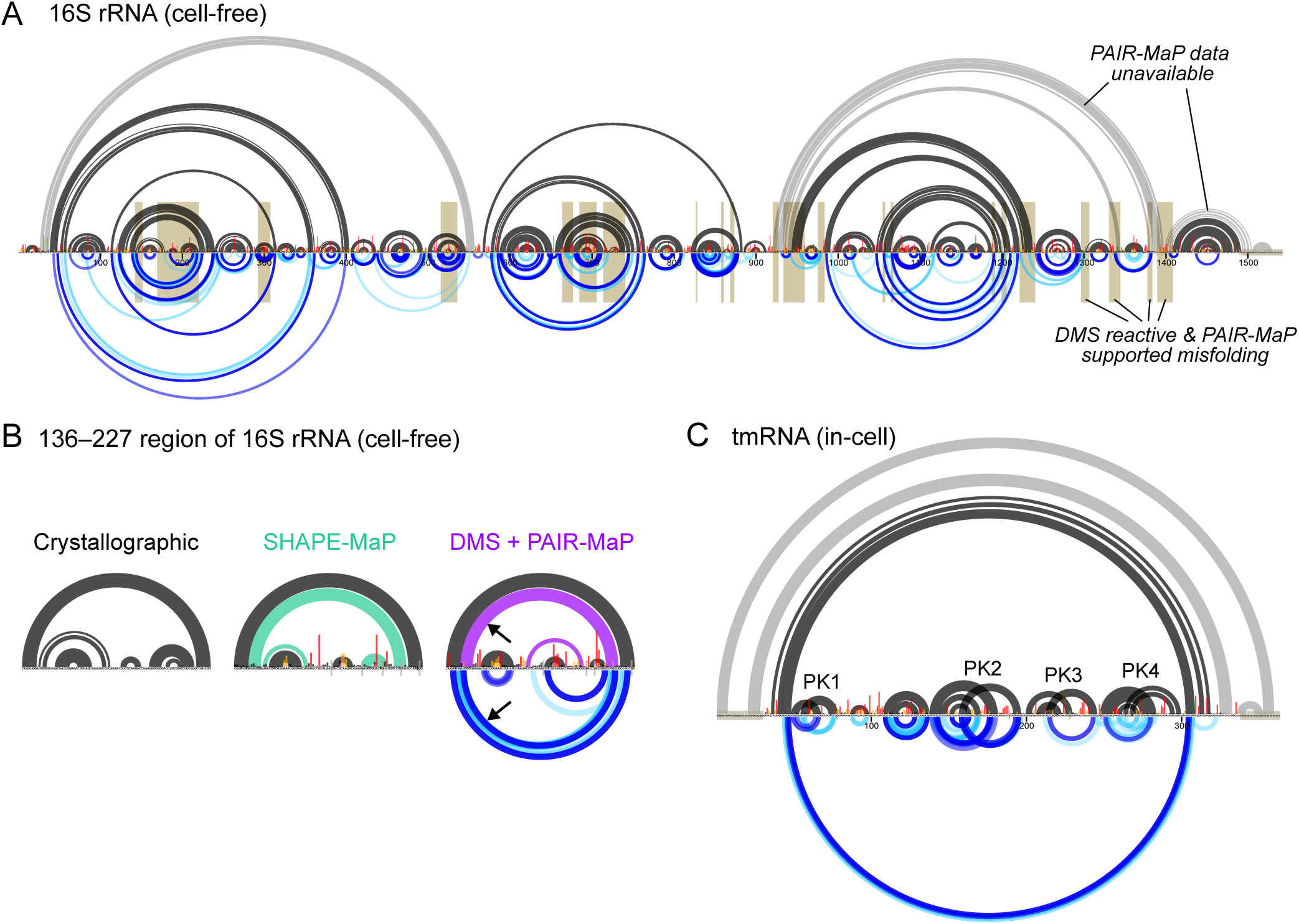
Detection of long-range base pairs, pseudoknots, and misfolding in endogenous RNAs. (**A**) PAIR-MaP data collected on the *E. coli* 16S rRNA under cell-free conditions. Misfolded regions indicated by high DMS reactivity and non-native PAIR-MaP signals are highlighted in gold. (**B**) Detail of misfolding in the 136-227 region of the 16S rRNA. Crystallographic, SHAPE-MaP (ref. (17)), and DMS+PAIR-MaP structure models are shown. PAIR-MaP correlations supporting the misfolded secondary structure are indicated by arrows. (**C**) In-cell PAIR-MaP data collected on *E. coli* tmRNA. Pseudoknots (PKs) are labeled. The key for PAIR-MaP plots is shown in Figure 2D.

Under cell-free conditions, principal PAIR-MaP correlations are highly predictive of the known secondary structure with an average positive predictive value (ppv) of 88%. Furthermore, many of the “false positive” correlations (corresponding to correlations that do not match the known secondary structure) are readily attributable to misfolding of the deproteinized RNAs and, indeed, provide the first direct evidence of such misfolding. Of particular note, we observe strong PAIR-MaP signals supporting misfolding of the 136–227 region of the 16S rRNA (Fig. 3B); prior SHAPE probing studies suggested that this region significantly populates an alternative conformation in the absence of proteins, but the validity of this misfolding event had remained controversial (8, 22, 23). We also observe PAIR-MaP signals supporting previously suggested misfolding events elsewhere in the 16S rRNA, 23S rRNA, and human RMRP (Fig. 3A, S3, S4) (8, 24, 25). When these regions with clear alternative folds are excluded, the average ppv of principal correlations increases to 92% (Table 1). The remaining false positives are likely a mixture of indirect interactions reflective of cooperative folding events and “real” misfolding interactions that we cannot confidently assess. Thus, we infer 92% represents a lower bound on the ppv of principal PAIR-MaP correlations.

**Table 1:** Accuracy of PAIR-MaP analysis and DMS+PAIR-MaP guided structure modeling. Positive predictive value (ppv) and sensitivity (sens) are reported for principal PAIR-MaP correlations, for complete structure models predicted using DMS+PAIR-MaP restraints, and for structure models predicted without experimental data (no data). Results are colored on a scale to highlight low (red) to high (green) accuracy. Note that model-free PAIR-MaP sensitivities are colored on a different scale, reflecting that 50% sensitivity is typically sufficient to define global RNA architecture. PKs, number of pseudoknots. RNAs known to have significant misfolding are colored gray and are excluded from averages.

|  |  |  | PAIR-MaP Principal Correlations |  | PAIR-MaP + DMS Model |  | No data Model |  |
| --- | --- | --- | --- | --- | --- | --- | --- | --- |
|  |  |  | ppv | sens | ppv | sens | ppv | sens |
| **Cell-free** | Length | PKs |  |  |  |  |  |  |
| 16S rRNA | 1542 | 0 | 81 | 41 | 79 | 85 | 46 | 51 |
| 16S (excl. misfold) | 1219 | 0 | 93 | 49 | 90 | 96 | 58 | 66 |
| 23S rRNA | 2904 | 0 | 84 | 37 | 79 | 85 | 53 | 54 |
| 23S (excl. misfold) | 2741 | 0 | 88 | 40 | 83 | 89 | 61 | 68 |
| 5S rRNA | 120 | 0 | 100 | 67 | 82 | 87 | 25 | 26 |
| RNase P | 377 | 2 | 81 | 35 | 87 | 87 | 64 | 67 |
| tmRNA | 363 | 4 | 88 | 37 | 88 | 90 | 63 | 58 |
| U1 snRNA | 164 | 0 | 100 | 43 | 88 | 98 | 88 | 98 |
| RMRP | 267 | 1 | 80 | 43 | 74 | 74 | 58 | 63 |
| **Average** |  |  | 92 | 45 | 86 | 91 | 60 | 64 |
| **In-cell** |  |  |  |  |  |  |  |  |
| tmRNA | 363 | 4 | 100 | 47 | 99 | 97 | 63 | 58 |
| U1 snRNA | 164 | 0 | 100 | 29 | 91 | 96 | 88 | 98 |
| RMRP | 267 | 1 | 92 | 64 | 97 | 97 | 58 | 63 |
| **Average** |  |  | 97 | 47 | 96 | 97 | 70 | 73 |

Minor PAIR-MaP correlations reveal additional complexities of non-coding RNA folding landscapes under cell-free conditions (Fig. 3, S2-S4). 30-50% of minor correlations correspond to native duplexes that are only partially folded under cell-free conditions. The other minor correlations are more challenging to evaluate. As noted above in our analysis of the adenine riboswitch, we expect some fraction of minor PAIR-MaP correlations to report indirect cooperative folding interactions. This is particularly evident for RNase P, where non-native PAIR-MaP signals are best explained as indirect interactions from global unfolding/folding transitions (Fig. S2). In contrast, for RNAs such as the 16S and 23S rRNA, a large fraction of the minor PAIR-MaP network almost certainly reflects alternative misfolded states (Fig. 3A, S4).

Strikingly, PAIR-MaP is also highly predictive of base-paired structure in cells for tmRNA, U1, and RMRP RNAs (ppv = 92−100% for principal correlations; Fig. 3C, S2, S3, Table 1). Indeed, principal and minor PAIR-MaP correlations markedly consolidate around the known structure of each RNA compared to cell-free PAIR-MaP networks, consistent with proteins stabilizing a single predominant structure in cells (Fig. S2, S3). However, PAIR-MaP analysis fails in cells for the rRNAs and RNase P. These RNAs are exceptionally stable such that paired nucleotides are almost never modified, such that PAIR-MaP cannot reliably measure and prioritize correlation signals. Such datasets are readily automatically identified and are rejected by the PAIR-MaP algorithm (SI Methods).

Overall, ∼45% of helices are detected as principal PAIR-MaP correlations (Table 1), but helix detection sensitivity (sens) does vary with molecular context. Analysis of our cell-free datasets reveals that PAIR-MaP has greatest sens (>50%) when each duplex strand contains an A or C (for example, AAG paired to CUU). Conversely, sens is lowest (<10%) when one strand consists entirely of G residues (GGG paired to CCC). This sequence dependence is consistent with the reactivity and specificity biases of DMS defined above. Sensitivity is additionally impacted by thermodynamic stability, with duplexes containing two or more G-C pairs detected with lower sens (for example, CCG paired to CGG is detected with ∼30% sens). As a single molecule method, PAIR-MaP also requires that duplexes occur in the same sequencing read, corresponding to an inter-duplex length limitation of ∼500 nts with current technology. Finally, sensitivity depends strongly on sequencing depth: a depth of at least ∼400,000 is required to reliably detect duplex correlations (Fig. S5).

In sum, PAIR-MaP is a specific and sensitive technique for directly detecting duplexes in endogenous RNAs in cells, revealing significant complexity in the folding landscapes of non-coding RNAs that is counteracted by protein stabilization in cells.

### Accurate in-cell structure modeling

While PAIR-MaP provides an important model-free strategy for detecting RNA duplexes and characterizing RNA structural complexity, a critical end-goal of chemical probing analysis is often to determine complete RNA structure models. Building on prior studies, we developed a strategy to use PAIR-MaP data to enable highly accurate RNA structure modeling, including in cells.

We first capitalized on our discovery that DMS reacts with all four nucleotides by developing new nucleotide-specific pseudo free energy change functions for DMS-directed structure modeling in *RNAstructure* (see SI Methods). On its own, this DMS-directed structure modeling strategy enables highly accurate *de novo* structure determination with average ppv ≈ 90% and sens ≈ 90% when applied to our panel of endogenous *E. coli* and human RNAs (Table S2). This level of accuracy is more than sufficient for mechanistic hypothesis generation. Nevertheless, some important structural features are missed, including one of the four pseudoknots in tmRNA (Fig. 4). We therefore developed an integrated modeling strategy in which we both apply per-nucleotide DMS reactivity restraints and also provide modest energetic bonuses to base pairs directly detected by PAIR-MaP. This integrated strategy is less accurate when modeling tmRNA under cell-free conditions due to the effects of several non-native PAIR-MaP correlations that likely reflect misfolding in non-cellular contexts (Table 1, S2). However, for all other RNAs and conditions, this integrated strategy yields equivalent or higher accuracy structure models (Table 1, S2). Notably, when using in-cell data, this integrated strategy recovers tmRNA structure with near perfect accuracy, including all four pseudoknots (ppv=99% and sens=97%; Fig. 4). It is worth emphasizing that tmRNA, with its mixture of long-range interactions and multiple pseudoknots, is one of the most difficult structure modeling challenges of which we are aware. Thus, while DMS-directed structure modeling provides excellent accuracy, integrated modeling with PAIR-MaP data can provide notable improvement for RNAs with particularly challenging structures.

**Figure 4:**
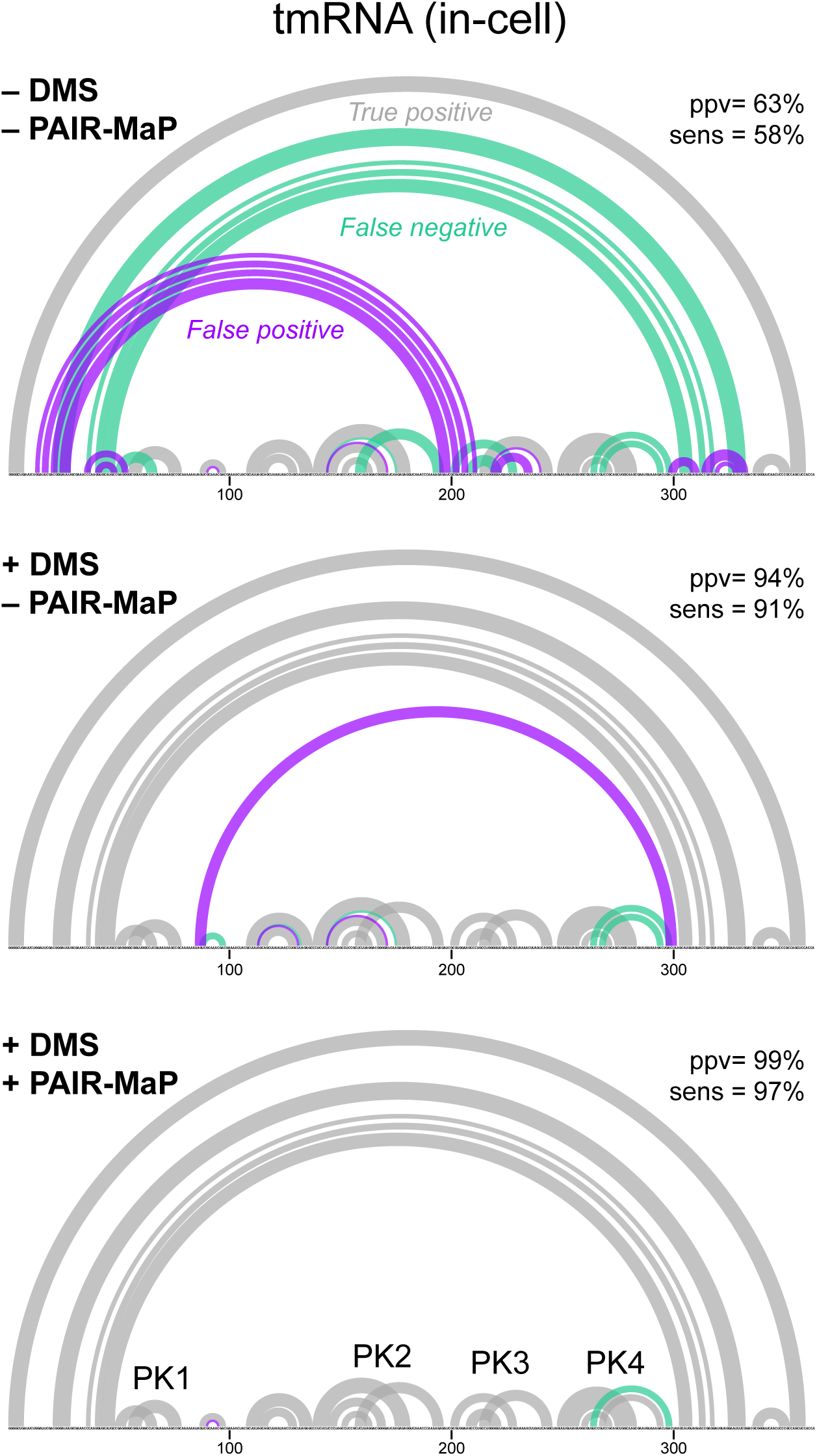
PAIR-MaP enables accurate modeling of tmRNA structure in cells. Secondary structure models are shown for modeling performed without experimental data (top), modeling guided by nucleotide-specific DMS reactivity restraints (middle), and DMS reactivity restraints and PAIR-MaP restraints (bottom). The four correctly modeled pseudoknots (PKs) are labeled.

Overall, the ∼90% accuracy of DMS+PAIR-MaP directed structure modeling is comparable to best-in-class SHAPE-based strategies (Table S2) (7). It is particularly striking that, using PAIR-MaP, accuracy remains similar or increases in cells for all RNAs. This analysis provides the first validation that DMS can be used to guide accurate structure modeling of long, complex RNAs. Furthermore, to our knowledge, this analysis provides the first general validation for any reagent that structure modeling can be performed accurately in cells.

### Identification of a novel conserved helix in RMRP

During benchmarking of PAIR-MaP on human RMRP it became clear that the accepted RMRP structure was incomplete. RMRP is an essential non-coding RNA, conserved across eukaryotes, that is involved in rRNA processing and other potential functions (26). RMRP is ancestrally related to the eukaryotic RNase P RNA, and phylogenetic analyses have shown that RMRP and RNase P share similar conserved structures (27, 28). As noted above, RMRP clearly misfolds under cell-free conditions (Fig. S3). This misfolding is resolved in cells, with PAIR-MaP correlations and structure modeling showing that RMRP forms the accepted base-paired structure with one notable exception. Strikingly, PAIR-MaP revealed the presence of an additional “P7” helix that closes the catalytic core domain of RMRP (Fig. 5A). The P7 helix is a conserved architectural feature of RNase P but, to date, has not been observed in RMRP (28). We used this updated structure model to realign the published RMRP multiple-sequence alignment (29), which newly reveals that the P7 extension is conserved from yeast to humans (Fig. 5B). Furthermore, single-nucleotide mutations or insertions within P7 are pathogenic in humans (Fig. 5B) (30). Thus, in-cell PAIR-MaP enabled discovery of a functionally important helix missed by previous analyses and provides new insights into the structural basis of human disease.

**Figure 5:**
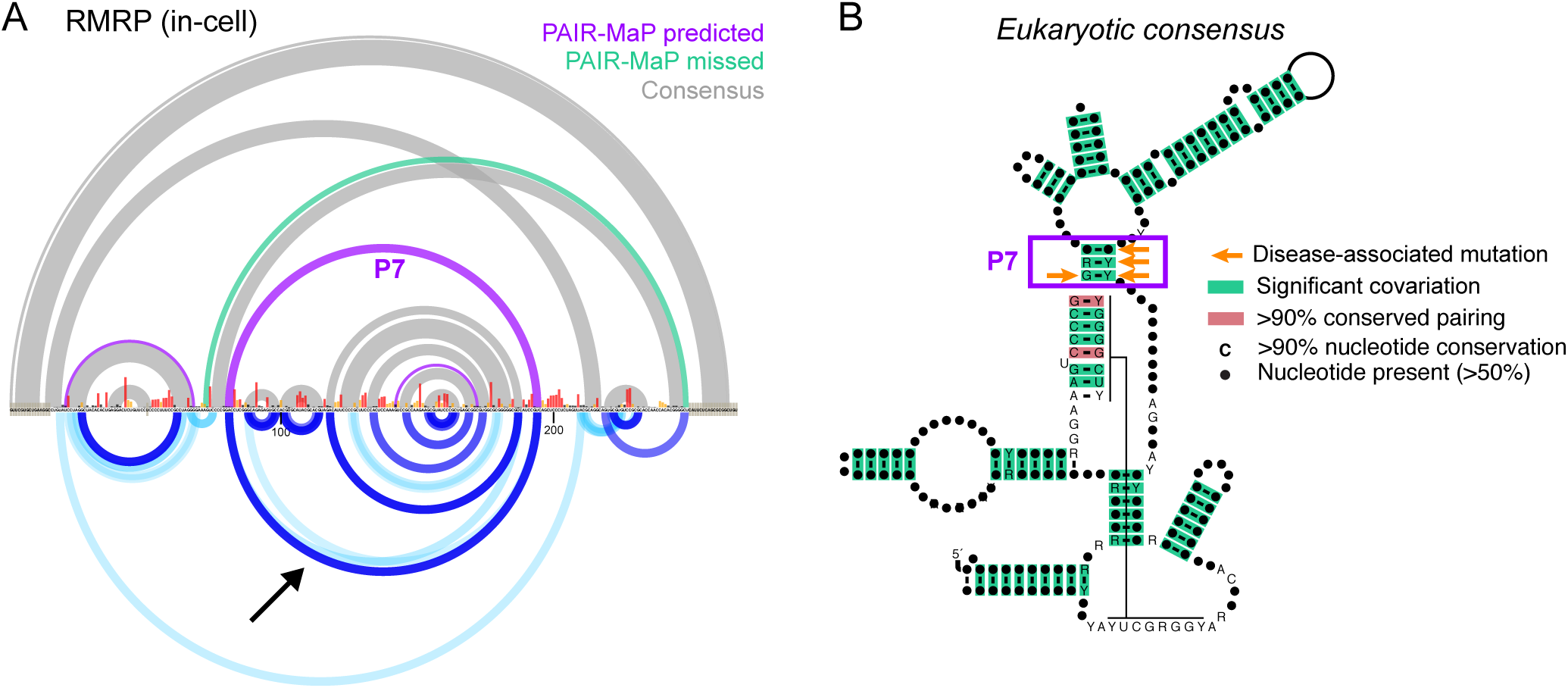
PAIR-MaP identifies a new conserved helix in RNase MRP. (**A**) Comparison between the in-cell DMS+PAIR-MaP structure model and prior covariation-based structure model (28). The PAIR-MaP correlation supporting the new P7 interaction is indicated by the arrow. The key for PAIR-MaP plots is provided in Figure 2D. (**B**) Realigned consensus structure of RMRP reveals significant covariation for the newly identified P7 helix (933 sequences, from yeast to human). Covariation was assessed using R-scape (39). Human disease-associated mutations in the P7 helix are indicated by orange arrows (30).

### Mechanistic insights into bacterial mRNA autoregulatory elements

We next applied PAIR-MaP to examine the structures of RNAs that have proven challenging to characterize via traditional approaches. Specifically, we focused on two *E. coli* 5' untranslated regions (5'-UTRs) that contain regulatory elements that bind ribosomal proteins (r-proteins) to inhibit translation of downstream genes, forming a feedback loop that ensures balanced synthesis of r-proteins and rRNA (31). These 5'-UTRs are good exemplars of the functional relationships between RNA structure and protein binding that govern regulation of many RNAs *in vivo*.

We first applied PAIR-MaP to characterize the S4-binding element (S4E) located upstream of the *E. coli rpsM* gene (also termed the α-operon). Prior studies have suggested that S4 induces a conformational change in the S4E, stabilizing a double pseudoknot structure that inhibits translation of *rpsM* and downstream genes (32, 33). However, the proposed double pseudoknot is not fully consistent with biochemical and genetic data, and the structure of the S4E in the absence of the S4 protein is unknown (SI Discussion; Fig. S6). Cell-free PAIR-MaP data show that the *rpsM* 5'-UTR and coding sequence (CDS) fold into four stem-loop helices (Fig. 6A). Of particular note, we identify a new, unstable helix (H3) formed between the Shine-Dalgarno sequence and the *rpsM* CDS. Strikingly, in-cell experiments reveal stabilization of H3 as well as appearance of new signals indicating loop-loop pairing between H2 and H3, consistent with S4 protein binding and stabilizing a kissing loop structure in cells (Fig. 6B). This kissing loop structure is more consistent with S4E sequence conservation compared to the previously proposed double pseudoknot, and uniquely explains the impact of Shine-Dalgarno sequence mutations on S4 binding (SI Discussion; Fig. 6B, C, S6) (31, 33). There is also potential structural homology between the kissing loop structure and the S4 binding site on the 16S rRNA (Fig. S6). Thus, our data support a model in which S4 binds a kissing loop structure, which stabilizes the H3 stem and thereby prevents translation initiation on *rpsM* (SI Discussion). We next examined the S2-binding element (S2E) located in the 5'-UTR of the *E. coli rpsB-tsf* transcript (34). Phylogenetic analyses have predicted that the S2E folds into a pseudoknot, but the pseudoknot interaction has not been directly confirmed and prior SHAPE experiments were ambiguous (31, 34, 35). Significantly, while no pseudoknot is observed under cell-free conditions, in-cell PAIR-MaP experiments reveal multiple minor signals consistent with S2-induced stabilization of the pseudoknot in cells (Fig. 6D,E). Our data also reveal a new “P1” helix (Fig. 6D,E). The functional importance of P1 is supported by prior genetic studies, which observed that deletion of P1-involved sequences abrogate S2 regulation (Fig. 6E) (34). The discovery of the P1 helix also illuminates how the S2 protein recognizes the S2E RNA. Whereas it was previously thought that the S2E lacked homology to the 16S rRNA binding site of S2 (31), identification of P1 makes it clear that S2 recognizes a common architecture in both the S2E and 16S rRNA (Fig. 6F). Interestingly, this architecture only appears to be conserved among enterobacterial S2Es, with S2Es from more distant bacterial species lacking the capacity to form P1 and thus potentially using a different mode of S2 recognition (SI Methods). Thus, in-cell PAIR-MaP analysis reveals divergent, yet functionally important, structural features of the *E. coli* S2E RNA that would be extraordinarily challenging to detect via conventional structure probing or covariation analyses.

**Figure 6:**
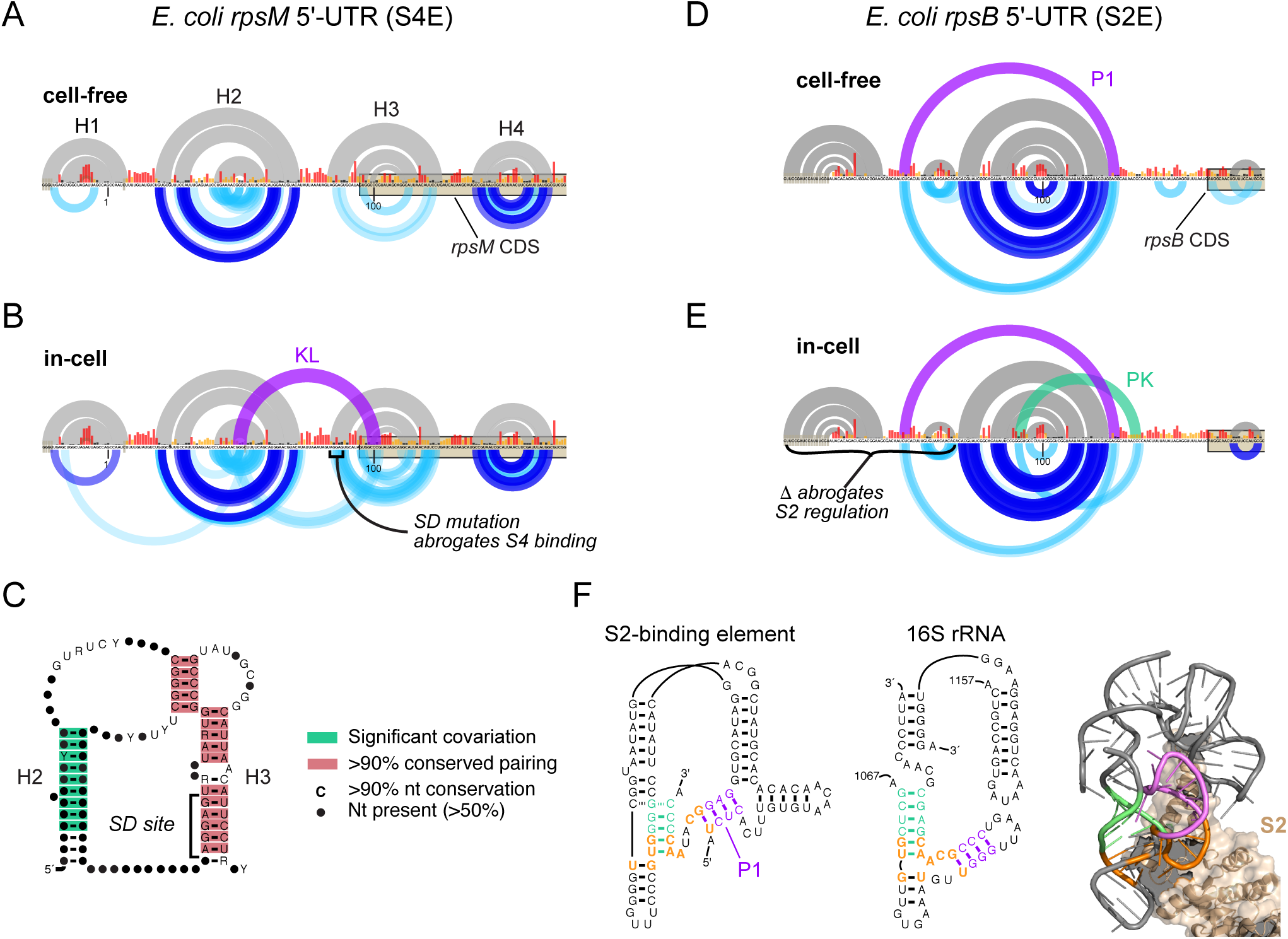
Novel structural features of r-protein regulatory elements. (**A**) Cell-free structure of the *E. coli rpsM* 5'-UTR containing the S4-binding element (S4E). Note that the sequence is numbered relative to the *rpsM*-specific transcription start site in accordance with prior studies; however, our data are specific to the intergenic form of the *rpsM* 5'-UTR transcribed from upstream promoters. (**B**) In-cell structure of the S4E. The kissing loop interaction (KL; purple) is not predicted by minimum free energy structure modeling but is clearly supported by PAIR-MaP correlations. (**C**) Revised consensus structure of the S4E across Gammaproteobacteria. Covariation and base-pairing conservation were assessed using R-scape and R2R, respectively (39, 41). (**D**) Cell-free structure of the *E. coli rpsB* 5'-UTR containing the S2-binding element (S2E). (**E**) In-cell structure of the S2E. The P1 and pseudoknot (PK) interactions (in purple and green, respectively) are not predicted by minimum free energy modeling but are clearly supported by PAIR-MaP correlations. (**F**) Homology between the S2E and the S2 binding site in the 16S rRNA. Homologous nucleotides are highlighted in orange. At right is the crystal structure of the S2 ribosome binding site (PDB: 4YBB). The key for PAIR-MaP plots is provided in Figure 2D.

## DISCUSSION

Chemical probing is a central tool in RNA structural biology, providing nucleotide-resolution insight into local RNA structure in a highly adaptable, experimentally concise manner. However, the inability to directly measure base pairing interactions had remained a fundamental limitation of these approaches. We show that single-molecule correlated chemical probing coupled with PAIR-MaP analysis resolves this limitation, allowing direct visualization of RNA duplexes in living bacterial and human cells. The correlation data obtained from PAIR-MaP experiments are typically sufficiently dense to define global RNA architecture, providing direct evidence of complex structural features such as pseudoknots and long-range pairing. Equally valuable, PAIR-MaP data provide insight into the complexity of the RNA structural landscape, revealing alternative and unstable pairing interactions that are exceptionally difficult to measure via conventional means. Finally, PAIR-MaP data can be used in combination with automated computational modeling strategies to derive complete, highly accurate models of RNA structure as it exists in cells.

PAIR-MaP offers major advantages compared to alternative *in vivo* duplex-detection strategies. Most importantly, PAIR-MaP resolves base pairs at nucleotide resolution with superior ppv (>90%) and sens (∼45%) (Table 1). PAIR-MaP is also simple and straightforward to implement; the innovation of the strategy lies in improved conditions enabling pan-RNA modification by DMS, the MaP readout, and algorithmic interpretation of the single-molecule correlated chemical probing signal. DMS-MaP experiments are already broadly used throughout the RNA community, and focused sequencing libraries of even rare RNAs can be easily prepared using PCR amplification without the need to enrich for or pull down target RNAs (36). Finally, a single PAIR-MaP experiment reports both local reactivity and pairwise interaction information, obviating the need for multiple experiments.

PAIR-MaP does have several limitations. In contrast to crosslinking and ligation strategies, PAIR-MaP requires duplexes to be self-contained within a contiguous sequencing read, currently ∼500 nucleotides, and cannot detect inter-molecular duplexes. PAIR-MaP also cannot detect duplexes in a few exceptionally stable, protein-coated RNP complexes such as the ribosome and RNase P. More generally, PAIR-MaP correlations are innately “non-native” measurements — in a sense, PAIR-MaP measures DMS-induced sequential unfolding of RNA molecules. The progressive accumulation of DMS adducts could promote formation of misfolded states, or shift the native equilibrium of dynamic RNAs. However, because of the stochasticity of the DMS modification process, every molecule is perturbed in a unique and non-coherent manner, and hence perturbations should average out over a population of molecules. Our extensive benchmarking supports that DMS-induced perturbations do not significantly impact PAIR-MaP accuracy.

Overall, our study highlights the extensive potential of PAIR-MaP for characterizing RNA structure and dynamics and ultimately for understanding biology. PAIR-MaP allowed us to determine the in-cell structure of human RMRP, revealing a new universally conserved and disease-linked RNA helix that has been missed by prior phylogenetic and chemical probing analyses. Our analysis of bacterial mRNA regulatory motifs further uncovered dynamic helices that are essential for understanding protein-binding and regulatory function, but which are only appreciably formed in cells and are invisible to lower-resolution methods. We anticipate that PAIR-MaP will broadly facilitate the next generation of high-resolution, in-cell structural insights into the many RNAs that continue to defy conventional characterization.

## MATERIALS AND METHODS

DMS probing experiments were performed on total RNA gently extracted (8, 37) from *E. coli* K-12 MG1655 and human Jurkat cells (cell-free) and on intact cells (in-cell) buffered with 200 or 300 mM bicine (pH 8.0), 200 mM potassium acetate (pH 8.0), and 5 mM MgCl_2_ at 37 °C. *In vitro* transcribed adenine riboswitch RNA was probed at 30 °C in the absence or presence of 100 μM adenine ligand in 300 mM bicine (pH 8.0), 100 mM NaCl, and 5 mM MgCl_2_. MaP reverse transcription was used to convert DMS adducts into mutations in cDNA, and is compatible with nearly all protocols for creating libraries for massively parallel sequencing (14, 36). Here, sequencing libraries were prepared by both randomly-primed Nextera (*E. coli* 16S and 23S rRNA) and gene-specific PCR (other RNAs) (36) and sequenced on an Illumina MiSeq instrument. *ShapeMapper* was used to align and parse mutations from DMS-MaP sequencing data (17). PAIR-MaP correlation analysis was performed using the newly developed *RingMapper*/*PairMapper* software suite. Correlations were computed between all pairs of 3-nt windows using the average product corrected G-test (38, 39) and then filtered by sequence complementarity and correlation strength. Structure modeling was performed using *RNAstructure* (40), incorporating DMS reactivities as nucleotide-specific free energy penalties and PAIR-MaP correlations as base-pair-specific energy bonuses. Detailed descriptions of these methods are provided in SI Appendix, Methods. The *RingMapper*/*PairMapper* software is available for download at https://github.com/Weeks-UNC/RingMapper.

## ACKNOWLEDGEMENTS

We thank S. Busan for helpful discussions and ongoing development of the *ShapeMapper* software, C. Weidmann for helpful discussions, and the Weeks lab for testing and feedback on PAIR-MaP. We thank D. Mathews (U. Rochester) for sharing access to the *RNAstructure* source code. This work was supported by the Arnold and Mabel Beckman Foundation (postdoctoral fellowship to A.M.M.) and the National Institutes of Health (R35 GM122532 to K.M.W.).

## DISCLOSURE

K.M.W. is an advisor to and holds equity in Ribometrix, to which correlated chemical probing technologies have been licensed.

## AUTHOR CONTRIBUTIONS

A.M.M. and K.M.W. designed research; A.M.M. performed experiments with assistance from P.S.I. and S.W.O.; A.M.M., N.L. and P.S.I. developed and performed analyses; A.M.M. and K.M.W. wrote the paper with input from all authors.

