## Supplementary material for "RNA base pairing complexity in living cells visualized by correlated chemical probing": SI Appendix

### SUPPORTING DISCUSSION

#### Prior experiments on the S4-binding element

The original double pseudoknot (DPK) model of the S4-binding element (S4E) was developed from early experiments mapping the impact of mutations on S4 binding (Fig. S6) (1). Following these initial experiments, the same library of S4E mutants has been extensively characterized by complementary biophysical and functional experiments (2-4). These data have been interpreted through the lens of the DPK model. However, as detailed below, much of this data is ambiguous and in several cases is inconsistent with the DPK structure.

The DPK structure is motivated by two pairs of mutations that disrupt and rescue S4 binding in filter-binding assays (1). Disruption of the proposed PK2 interaction by mutation 17 reduces S4 binding ~10-fold and is rescued by the compensatory mutation 21 (Fig. S6A). Disruption of PK3 by mutation 18 reduces S4 binding ~2-fold and is rescued by the compensatory mutation 22. However, the significance of these compensatory mutations is less clear when considered in the context of other mutation data. Relative to mutations spanning the kissing loop (KL) structure that uniformly and catastrophically disrupt binding, mutations to the PK2 and PK3 regions have inconsistent and modest impact on binding (Fig. S6A). Notably, multiple mutations that should disrupt PK2 and PK3 have no impact on binding. Subsequent *in vivo* functional assays also are inconsistent with the DPK model (2). Significantly, what should be compensatory 18+22 mutations do not rescue S4 repressive function *in vivo*. Finally, biophysical characterization of different S4E mutants also reveals inconsistencies of the DPK model, with the expected compensatory mutations (17+21 and 18+22) failing to rescue S4E tertiary folding (3). Overall, the inconsistent and modest impact of mutations to the proposed PK2 and PK3 interactions and the inconsistency of compensatory rescue argue against direct PK2 and PK3 pairing. Rather, these data are more consistent with these regions interacting indirectly, as would be expected for a kissing-loop type structure where nucleotides adjacent to the KL duplex contribute to tertiary stability and S4 binding but do not directly pair (Fig. S6D).

While most of the mutations tested by prior studies are not expected to impact H3, three mutations do provide evidence supporting H3 and the kissing-loop structure. Mutation 19 (G95→A) converts a G•U pair in H3 to an A•U pair, and as expected, has no impact on S4 binding *in vitro* or repression function *in vivo* (2). Mutation 18 modestly disrupts H3 and modestly decreases S4 binding affinity, although has no impact on repression function *in vivo* (1). Most significantly, mutation of the AGGAG Shine-Dalgarno sequence (SD; Fig. S6A) to its sequence complement, UCCUC, completely disrupts H3 and, as expected, completely abolishes S4 binding (4). By comparison, in the context of the DPK model, the SD mutation occurs in the middle of a 25-nt single-stranded loop and thus would not be expected to have such profound impact.

Considered as a whole, the body of mutational, phylogenetic, and structural data from past studies and our current study strongly support the kissing-loop structure as the structure recognized by S4.

**Implications for mechanism S4 translation inhibition**

R-protein translation autoregulatory elements have historically been proposed to function via one of two distinct mechanisms: (i) a displacement mechanism where r-protein binding prevents (or displaces) the 30S subunit from binding the mRNA, or (ii) an entrapment mechanism where the r-protein•mRNA complex binds the 30S subunit but is then trapped in an inactive translation-incompetent state (5). Notably, the S4E RNA has served as the primary example of the entrapment mechanism (4-6). The S4•S4E binary complex binds the 30S subunit, forming a ternary complex that is translationally inactive (4). In the double pseudoknot model of the S4•S4E complex, the SD sequence is located in a long single-stranded loop and was thus proposed to mediate ternary complex formation via SD•antiSD pairing. However, the SD sequence is internally base paired in our kissing-loop model, preventing SD•antiSD pairing. While the SD•antiSD interaction has traditionally been viewed as critical to mRNA binding to 30S subunits, recent studies have emphasized that 30S subunits interact with single-stranded A/U-rich elements such as found between H2 and H3 in the S4E (Fig. 6B, S6D) (7, 8). Thus, the kissing-loop structure does not preclude formation of an entrapped S4•S4E•mRNA ternary complex. Nevertheless, we suggest a modest revision of the mechanism of S4 translation inhibition that builds on the renewed appreciation of RNA unfolding kinetics as a major rate-limiting step of translation initiation (9). In the absence of S4, the H3 helix (which overlaps the SD sequence and start codon of *rpsM*) is very unstable, allowing efficient unfolding and accommodation of the *rpsM* mRNA into the 30S mRNA cleft. However, upon binding by the S4 protein, H3 and the KL interaction are stabilized and the energy barrier to S4E unfolding is significantly increased. Thus, in the kinetic competition between accommodation of the *rpsM* mRNA versus disassociation from the 30S subunit, S4 binding favors disassociation, thereby inhibiting translation. (Note that observations of stable S4•S4E•mRNA ternary complexes were made at low-temperature, non-physiological conditions (4, 6); we expect that S4•S4E•mRNA complexes are only transiently stable at physiological conditions.)

### SUPPORTING METHODS

#### Identification of alkaline, non-quenching DMS buffer conditions

Most biological buffering reagents, including HEPES, react directly with DMS and will partially quench the DMS modification reaction when present at the concentrations necessary to appropriately buffer the reaction (10). We identified bicine [2-(bis(2-hydroxyethyl)amino)acetic acid] as a buffering reagent that can both maintain a moderately basic pH and allow significant RNA modification. Depending on reaction conditions (including the presence of other buffering salts such as potassium acetate), 200 to 300 mM bicine (pH = 8.0) is sufficient to maintain pH control throughout the reaction (final pH ≈ 7, after 6-10 min reaction). Note that buffer stocks prepared at room temperature should be titrated to pH 8.3 and will then have pH 8.0 at 37 °C due to the temperature dependence of the bicine $pK_a$. Bicine pHs listed below correspond to the value at 37 °C. We note that a prior study also observed that U residues are DMS labile at pH>8, but this property has not been previously harnessed for structure probing (11).

#### DMS probing of the adenine riboswitch

A DNA template of the *add* adenine riboswitch from *V. vulnificus* flanked by 5′ and 3′ structure cassettes (12) was synthesized (IDT; Table S3), amplified by PCR (Q5 DNA polymerase, NEB), and purified (PureLink PCR column, Invitrogen). RNA was transcribed *in vitro* [400 µL; 40 mM Tris (pH 8.0), 25 mM $MgCl_2$, 2.5 mM Spermidine, 0.01% (vol/vol) Triton X-100, 10 mM DTT, 5 mM each NTP, ~5 µg DNA template, 0.05 mg/mL T7 RNA polymerase (lab made), 0.5 U/mL pyrophosphatase (NEB); 37 °C; 4h], purified (Agencourt RNAclean XP beads; Beckman

Coulter), and stored at −20 °C. RNA size and purity was confirmed using Bioanalyzer analysis and concentration was quantified (Qubit RNA BR assay, Invitrogen).

For probing experiments, RNA [3 pmol in 3 μL 0.5× TE (pH 8.0)] was first denatured at 95 °C for 2 min followed by snap cooling on ice for 2 min. RNA was folded by adding 4 μL of 2.5× folding buffer [750 mM bicine (pH 8.0), 250 mM NaCl, 12.5 mM $MgCl_2$] and incubating at 30 °C for 10 min, and then adding either 2 μL 500 μM adenine or $H_2O$ and incubating at 30 °C for an additional 20 min. 9 μL folded RNA was added to 1 μL DMS solution (1.7 M in EtOH), allowed to react for 10 min at 30 °C, and then quenched using 10 μL neat 2-mercaptoethanol (2ME) and placed on ice. Note that the 30 °C folding and modification temperature was chosen to match conditions used in prior NMR experiments (13), and the 10 min reaction time was chosen to achieve a comparable level of DMS modification as standard 37 °C reaction conditions. Following quenching, RNA was precipitated with ethanol and resuspended in $H_2O$. No-reagent control samples were treated identically, substituting neat EtOH for the DMS solution.

**DMS probing of *E. coli* total RNA**

***Cell-free experiments.*** 2 mL overnight culture of *E. coli* K-12 MG1655 was used to inoculate 148 mL LB and grown to $OD_{600}$≈0.5 at 37 °C under vigorous shaking. 16.65 mL of 187.5 μg/mL rifampicin was added and the culture incubated for 20 min at 37 °C under shaking to chase ribosomes to fully assembled states (14). Total RNA was then gently extracted following a previously published protocol with slight modifications (15). 48 mL of culture was pelleted at 14,000*g* at 4 °C for 7 min. Cell pellets were resuspended in 32 mL lysis buffer [15 mM Tris (pH 8.0), 450 mM sucrose, 8 mM EDTA], 1.28 mL lysozyme (10 mg/mL) was added, and the solution incubated at 23 °C for 5 min and on ice for 10 min. Protoplasts were collected by centrifugation at 5,000*g* at 4 °C for 5 min, resuspended in 3.8 mL protoplast lysis buffer [50 mM HEPES (pH 8.0), 200 mM NaCl, 5 mM $MgCl_2$, 1.5% (wt/vol) SDS], followed by addition of 76 μL Proteinase K (10 mg/mL) and incubated at 23 °C for 5 min and then on ice for 10 min. SDS was precipitated by adding 1 mL precipitation buffer [50 mM HEPES (pH 8.0), 1M KOAc, 5 mM $MgCl_2$] and centrifugation at 14,000*g* for 7 min. The RNA-containing eluent was extracted 3 times using phenol/chloroform/isoamyl alcohol (PCA) that was pre-equilibrated with 1×

folding buffer [200 mM bicine (pH 8.0), 200 mM KOAc, 5 mM $MgCl_2$], and 3 times with chloroform. RNA was exchanged into 1× folding buffer (PD-10 columns, GE Healthcare). RNA in folding buffer was equilibrated at 37 °C for 6 min, and then 9 volumes of RNA solution were added to 1 volume of DMS solution (1.7 M DMS in EtOH) and reacted for 6 min at 37 °C. Reactions were quenched via addition of 10 volumes ice-cold 20% 2ME [in $H_2O$ (vol/vol)] followed by ethanol precipitation. Contaminating genomic DNA was removed (TURBO DNase, Invitrogen) incubating at 37 °C for 60 min (0.02 U/μL additional enzyme was spiked-in after the initial manufacturer recommended 30 min incubation), followed by purification (RNeasy MinElute columns, Qiagen). Concentrations were measured by Qubit RNA BR assay. No-reagent control samples were treated identically, substituting neat EtOH for the DMS solution.

***In-cell experiments.*** Initial experiments on intact *E. coli* cells revealed that standard DMS modification conditions (170 mM DMS final concentration) yielded insufficient RNA modification for PAIR-MaP analysis. In-cell *E. coli* probing experiments were thus performed using increased DMS concentrations (4× relative to standard) with corresponding increased buffer concentrations. 49 mL LB was inoculated with 1 mL overnight culture of *E. coli* K-12 MG1655 and grown to $OD_{600}$≈0.5 at 37 °C under vigorous shaking. 5.55 mL of 187.5 μg/mL rifampicin was added and the culture incubated at 37 °C with shaking for 20 min. 45 mL of culture was pelleted at 3,200*g* for 5 min and resuspended in 20 mL folding buffer [300 mM bicine (pH 8.0), 200 mM KOAc, 5 mM $MgCl_2$] and equilibrated at 37 °C for 5 min. 4.5 mL buffered cells were transferred to a new tube containing 500 μL 4× DMS solution (6.8 M DMS in EtOH), mixed vigorously, and incubated for 6 min at 37 °C. DMS reactions were quenched by adding 20 mL ice-cold 20% 2ME and placed on ice. Cells were pelleted at 5,000*g* for 7 min at 4 °C, resuspended in 1 mL lysozyme solution [1 mg/mL lysozyme in 0.5× TE (pH 8.0)], and incubated on ice for 5 min. Total RNA was extracted (using 8 mL TRIzol reagent, Invitrogen) followed by isopropanol precipitation. RNA was treated with TURBO DNase using the same procedure described above and purified (Agencourt RNAclean XP beads, Beckman Coulter). Bioanalyzer analysis was used to confirm RNA integrity (RIN>8.0) and concentration was determined using the Qubit RNA BR assay. No-reagent control samples were treated identically, substituting neat EtOH for the DMS solution.

**DMS probing of total human RNA**

***Cell culture.*** Jurkat cell cultures were maintained in RPMI 1640 media (Gibco) supplemented with 10% FBS, 50 U/mL penicillin, and 50 μg/mL streptomycin with 5% $CO_2$ at 37 °C. Two days prior to probing experiments, 1×$10^6$ cells were transferred to 30 mL fresh media in a T75 culture flask. Immediately prior to the probing experiment, cells were spun down at 1,000*g* for 3 min, washed with DPBS, and resuspended in 1 mL fresh media.

***Cell-free experiments.*** Cell nuclei were isolated by pelleting 5×$10^6$ cells at 1,000*g*, resuspending in lysis buffer [40 mM Tris (pH 8.0), 25 mM NaCl, 6 mM $MgCl_2$, 1 mM $CaCl_2$, 256 mM Sucrose, 0.5% Triton X-100, 0.45 U/μL DNase I (Roche), 0.5 U/μL RNasin (Promega)] and incubating at 4 °C for 5 min. Nuclei were then pelleted at 4 °C for 2 min at 2,250*g*, resuspended in Proteinase K buffer [40 mM Tris (pH 8.0), 200 mM NaCl, 1.5% SDS, 0.5 μg/μL Proteinase K], and incubated at 25 °C for 45 min. RNA was extracted using phenol/chloroform/isoamyl alcohol pre-equilibrated with 1× folding buffer [200 mM bicine (pH 8.0), 200 mM KOAc, 5 mM $MgCl_2$] following previously described procedures (16). RNA was exchanged into 1.1× folding buffer (PD-10 columns, GE Healthcare) and incubated at 37 °C for 20 min. 9 volumes of RNA were then added to 1 volume of DMS solution (1.7 M DMS in EtOH), reacted for 6 min at 37 °C, and quenched by addition of 10 volumes of ice cold 20% 2ME. Reactions were purified using isopropanol precipitation, followed by treatment with TURBO DNase, and purified (Mag-Bind TotalPure NGS beads, Omega). RNA concentrations were determined using the Qubit RNA HS Assay (Invitrogen). No-reagent control RNA was prepared identically, substituting neat EtOH for the DMS solution.

***In-cell experiments.*** Cultures containing 5×$10^6$ cells in fresh media were supplemented with 200 mM bicine (pH 8.0) buffer (final concentration). 9 volumes of buffered cell culture were added to 1 volume of DMS solution (1.7 M DMS in EtOH) and allowed to react for 6 min at 37 °C. Reactions were quenched by adding 10 volumes of ice cold 20% 2ME. Cells were pelleted at 1,000*g* for 3 min followed by isolation of total RNA (using 1 mL of TRIzol reagent). Contaminating genomic DNA was removed (TURBO DNase, Invitrogen) and RNA purified (Mag-Bind TotalPure NGS beads, Omega). RNA concentrations were determined using the

Qubit RNA HS Assay. No-reagent control RNA was prepared identically, substituting neat EtOH for the DMS solution.

**Reverse transcription**

Mutational profiling (MaP) reverse transcription (RT) was performed using an updated protocol that substantially improves yield and product length (G.M. Rice and K.M. Weeks, unpublished). Note that MaP is compatible with nearly all downstream methods for preparing libraries for massively parallel sequencing; here, we used both random-primed (total RNA) and gene-specific strategies. In a 10 µL volume, RNA was mixed with 200 nM specific primer (or 20 ng/µL random 9mer) (Table S3) and 2 mM dNTPs and incubated at 65 °C for 10 min followed by 4 °C for 2 min. 9 uL 2.22× MaP buffer [1× MaP buffer consists of 6 mM $MnCl_2$, 1 M betaine, 50 mM Tris (pH 8.0), 75 mM KCl, 10 mM DTT] was added and the combined solution incubated at 23 °C for 2 min. Finally, 1 µL SuperScript II Reverse Transcriptase (Invitrogen) was added and the RT reaction was performed according to the following temperature program: 25 °C for 10 min, 42 °C for 90 min, 10×[50 °C for 2 min, 42 °C for 2 min], 72 °C for 10 min. For the adenine riboswitch, ~1 pmol RNA was input into RT. For *E. coli* and human short RNAs, 0.5-2 µg total RNA was input into RT. For randomly primed *E. coli* RNA samples (16S and 23S rRNA), 1.5-3 µg total RNA was input into RT. RT products were then purified (Agencourt RNAclean XP beads, Beckman Coulter).

**Library preparation and sequencing**

***Adenine riboswitch.*** Sequencing libraries for the adenine riboswitch were prepared from specifically primed cDNA products using the two-step PCR approach (17) (Table S3). 1 µL cDNA was input to PCR1, performed as follows: 98 °C for 8 s, 10 cycles of [98 °C for 8 s, 66 °C for 20 s, 72 °C for 20 s], and 72 °C for 2 min. PCR1 product was purified (PureLink PCR Micro columns, Invitrogen). ~5 ng PCR1 product was input to PCR2, which was performed as follows: 98 °C for 30 s, then 10 cycles of [98 °C for 8 s, 68 °C for 20 s, 72 °C for 20 s], and 72 °C for 2 min. PCR2 products were purified (Agencourt AMPure XP beads; 0.8× bead ratio), and sequenced with an Illumina MiSeq instrument using 2×150 paired-end sequencing (v2 chemistry).

***Small RNAs.*** Sequencing libraries for the *E. coli* 5S rRNA, tmRNA, RNase P, S2-binding element, and S4-binding element, and human RMRP and U1 snRNA were prepared from specifically primed cDNA products using the two-step PCR approach (17) (Table S3). 1–2.5 µL cDNA was input to PCR1, which was performed as follows: 98 °C for 30 s, 18 cycles of [98 °C for 10 s, 60 °C for 30 s, 72 °C for 20 s], and 72 °C for 2 min. PCR1 product was purified (Agencourt AMPure XP beads; 0.7× bead ratio for RNase P and tmRNA, 1.2× bead ratio for other RNAs). 0.5–1 ng PCR1 product was input to PCR2, performed as follows: 98 °C for 30 s, then 12 cycles of [98 °C for 10 s, 66 °C for 30 s, 72 °C for 20 s], and 72 °C for 2 min. PCR2 products were purified (Agencourt AMPure XP beads; 0.7× bead ratio for RNase P and tmRNA, 1.2× bead ratio for other RNAs) and sequenced with an Illumina MiSeq instrument. 2×300 sequencing (v3 chemistry) was used for RNase P and tmRNA libraries and 2×150 sequencing (v2 chemistry) was used for the remaining libraries.

***Total E. coli RNA.*** Libraries of total RNA (16S and 23S rRNA datasets) were prepared using the randomer Nextera workflow (17). Purified randomly primed cDNA product was converted to double-stranded DNA using the NEBNext second-strand synthesis module (NEB) with a 150 min incubation at 16 °C. The resulting DNA was purified and size-selected (Agencourt AMPure XP beads; 0.65× bead ratio). Nextera XT (Illumina) was used to construct libraries according to the manufacturer's protocol, followed by purification and size-selection (Agencourt AMPure XP beads; 0.56× bead ratio). Bioanalyzer analysis indicated average library sizes of 478 bp for cell-free modified RNA, 823 bp for cell-free untreated RNA, 800 bp for in-cell modified RNA, and 930 bp for in-cell untreated RNA. Libraries were sequenced using 2×300 paired-end sequencing on an Illumina MiSeq instrument (v3 chemistry).

**Sequence alignment and mutation parsing**

*ShapeMapper* (v2.1.4) was used to align and parse mutations from DMS-MaP sequencing data (18). The *--amplicon* option was used for amplicon datasets, which masks out primer regions and filters out spurious reads that do not align at the expected 5′ and 3′ primer sites. The *--random-primer-len 9* option was used to mask RT primer sites in randomly primed datasets (16S and 23S rRNAs). The *--output-parsed* option was used to generate parsed mutation files, which parse each aligned read into two binary strings: a "depth" string indicating coverage at each nucleotide position (e.g. the base call meets the Phred score threshold), and a "mutation" string indicating valid mutation events (modifications) called by *ShapeMapper*. These parsed mutation files serve as the inputs to *RingMapper*/*PairMapper* correlation analysis (see below). Default parameters were used for all other *ShapeMapper* settings. Initial analysis of human U1 snRNA datasets revealed the presence of two minor sequence variants. Therefore, to specifically segregate reads from the primary U1 species, U1 snRNA alignments were done against three reference sequences (the primary and two inferred minor variants). The two minor variants accounted for <5% of total reads. All reference sequences used for alignments are provided in supporting data.

**PAIR-MaP analysis**

***Software.*** We created a new suite of software tools for computing and plotting correlated modifications observed in single-molecule chemical probing experiments. *RingMapper* is a fast, general purpose analysis code for calculating statistically significant correlations from pre-processed parsed-mutation files output by *ShapeMapper. PairMapper* uses *RingMapper* as a backend to compute correlations, and then filters and prioritizes the correlations to identify principal and minor PAIR-MaP signals. *arcPlot* is a flexible plotting tool that can plot *RingMapper* and *PairMapper* correlation outputs as well as RNA structure files (.ct format) and DMS reactivity profiles, and was used to generate all structure/correlation figures. A complete description of *arcPlot* will be provided in a future publication (Mustoe et al., *unpublished*). The software is written in a mixture of Python (2.7) and Cython and is available for download with documentation at https://github.com/Weeks-UNC.

***Correlation measurement.*** In PAIR-MaP analysis, correlations are evaluated between all pairs of three-nucleotide windows. The state of each nucleotide window, $n_{[i,\, i+2]}$, is assigned as ***u*** (unmodified), ***m*** (modified), or ***nd*** (no data):

$$n_{[i,i+2]} = \begin{cases} \boldsymbol{u} \quad if \left[\sum_{k=i}^{i+2} \mathbb{D}(k)\right] \geq 1 \ \ \&\& \ \left[\sum_{k=i}^{i+2} \mathbb{M}(k)\right] == 0 \\ \boldsymbol{m} \quad if \left[\sum_{k=i}^{i+2} \mathbb{D}(k)\right] \geq 1 \ \ \&\& \ \left[\sum_{k=i}^{i+2} \mathbb{M}(k)\right] \geq 1 \\ \boldsymbol{nd} \ \ if \left[\sum_{k=i}^{i+2} \mathbb{D}(k)\right] == 0 \end{cases}$$

where $\mathbb{D}$ is the "depth string" and $\mathbb{M}$ is the "mutation string" for an aligned read parsed by *ShapeMapper*. In other words, if at least one position of the window is covered by the read and there are no mutations then the window is considered unmodified. If there is at least one mutation then the window is considered modified. For each pair of windows there are four possible states, [($\boldsymbol{u}$, $\boldsymbol{u}$), ($\boldsymbol{u}$, $\boldsymbol{m}$), ($\boldsymbol{m}$, $\boldsymbol{u}$), and ($\boldsymbol{m}$, $\boldsymbol{m}$)], forming a 2×2 contingency table that is tabulated over all reads (***nd*** values are skipped).

Statistical correlation between windows is evaluated using the G-test (19). For two windows [*i*, *i*+2] and [*j*, *j*+2] (shorthand *i* and *j*) the uncorrected G statistic is

$$G_{ij} = 2\, N_{ij} \sum_{\substack{a \in \{\boldsymbol{u},\boldsymbol{m}\} \\ b \in \{\boldsymbol{u},\boldsymbol{m}\}}} p_{ij}(a,b) \ln\left(\frac{p_{ij}(a,b)}{p_i(a)p_j(b)}\right)$$

$N_{ij}$ is the total number of joint observations. $p_{ij}(a,b) = Obs_{ij}(a,b)/N_{ij}$, where $Obs_{ij}(a,b)$ is the observed counts of state *(a, b)*. $p_i(a)$ is the marginal frequency of state *a*, $p_i(a) = \left(Obs_{ij}(a,\boldsymbol{u}) + Obs_{ij}(a,\boldsymbol{m})\right)/N_{ij}$.

The uncorrected G statistic depends systematically on the average modification rate of each nucleotide window (20). This systematic dependence is normalized out by the average product correction (APC) of the G statistic (20, 21):

$$G_{ij}^{APC} = G_{ij} - \frac{\overline{G_i}\,\overline{G_j}}{\bar{G}}$$

$\overline{G_i}$ and $\overline{G_j}$ are the average G statistics of windows *i* and *j*,

$$\overline{G_i} = \frac{\sum_j G_{ij}\, I_A(i,j)}{\sum_j I_A(i,j)}$$

$\bar{G}$ is the average G statistic of all windows:

$$\bar{G} = \frac{\sum_{i,j} G_{ij}\, I_A(i,j)}{\sum_{i,j} I_A(i,j)}$$

$I_A(i,j)$ is the indicator function, equal to 1 for *(i, j)* pairs passing proximity, depth, and background filters (described below), and equal to 0 otherwise.

A pair of nucleotide windows is considered significantly correlated if $G_{ij}^{APC} > 20$, corresponding to $P < 10^{-5}$.

***Proximity, depth, and background filtering.*** Nucleotide windows separated by 8 or fewer nucleotides are excluded from analysis to filter out local read alignment artifacts ($I_A(i, j) = 0$ for $j \leq i+8$). Windows with insufficient observations are filtered out ($I_A(i, j) = 0$ if $N_{ij} < 10{,}000$ or $\min(Obs_{ij}) < 50$). The unmodified control dataset is also used to identify windows with high background mutation rates and statistically significant background correlations, which are deemed unreliable and filtered out. Specifically, $I_A(i, j) = 0$ for all $j$ if $r_{bg}(i) > 0.06$, where $r_{bg}(i)$ is the background mutation rate of window $i$, and $I_A(i, j) = 0$ if $G_{ij,bg} > 10.83$ (significant at P<0.001), where $G_{ij,bg}$ is the uncorrected G statistic computed for $(i, j)$ in the unmodified background sample. (Note that the $G_{ij,bg}$ filter is not applied if $N_{ij,bg} < 10{,}000$ or $\min(Obs_{ij,bg}) < 5$).

***PAIR-MaP filtering and prioritization.*** High-confidence base pairing signals are identified from the set of significantly correlated nucleotides ($G_{ij}^{APC} > 20$) by filtering by sequence complementarity, correlation strength, and reactivity (Fig. S1). Correlated nucleotide windows $i$ and $j$ must be able to form three Watson-Crick or G-U (WC/GU) pairs. $i$ and $j$ must be positively correlated, defined as $p_{ij}(\boldsymbol{m}, \boldsymbol{m}) > p_i(\boldsymbol{m})\, p_j(\boldsymbol{m})$, where $p_{ij}(\boldsymbol{m},\boldsymbol{m})$ is the joint frequency of the $(\boldsymbol{m}, \boldsymbol{m})$ state and $p_i(\boldsymbol{m})$ and $p_j(\boldsymbol{m})$ are the marginal frequencies. The $(i, j)$ correlation must also be two standard deviations greater than the average correlation observed at $i$ and $j$, defined as $z_{ij} \geq 2$ where

$$z_{ij} = \frac{1}{2}\left(\frac{G_{ij}^{APC} - \mu_i}{\sigma_i} + \frac{G_{ij}^{APC} - \mu_j}{\sigma_j}\right)$$

$\mu_i$ (and $\mu_j$ by symmetry) is the mean of $G_{ij}^{APC}$ averaged over all $j$:

$$\mu_i = \frac{\sum_j G_{ij}^{APC}\, I_A(i,j)}{\sum_j I_A(i,j)}$$

$\sigma_i$ (and $\sigma_j$ by symmetry) is the standard deviation of $G_{ij}^{APC}$ over all $j$:

$$\sigma_i = \left[\frac{\sum_j \left(G_{ij}^{APC} - \mu_i\right)^2 I_A(i,j)}{\sum_j I_A(i,j)}\right]^{1/2}$$

The complementary *(i, j)* correlations passing these filters are then prioritized as "principal" and "minor" correlations. Principal correlations comprise the strongest complementary correlation observed at both windows ($G_{ij}^{APC} \geq G_{ik}^{APC}$ and $G_{ij}^{APC} \geq G_{lj}^{APC}$ for all *k* and *l* in the set of complementary correlations) and additionally satisfy the constraint $\overline{r_i} < 0.2$ and $\overline{r_j} < 0.2$, where $\overline{r_i}$ is the mean normalized reactivity of window *i*. The remaining set of complementary correlations that satisfy $\overline{r_i} < 0.5$ and $\overline{r_j} < 0.5$ are classified as minor.

***Restraints for minimum free energy structure modeling.*** PAIR-MaP correlations are used to guide minimum free energy structure modeling by giving small energetic bonuses to PAIR-MaP-supported base-pairs (an approach adapted from prior studies (22, 23)). Whereas stringent filtering criteria are used to identify high-confidence base pairs in model-free PAIR-MaP analysis, filtering thresholds are relaxed for structure modeling applications, allowing the global free energy minimization process to resolve conflicting and/or non-specific PAIR-MaP signals. All correlated windows with $G_{ij}^{APC} > 20$ that can form at least two WC/GU pairs, are positively correlated (as defined above), and with $z_{ij} \geq 1$ (as defined above) are given energetic bonuses according to

$$\Delta G_{pm}\left(G_{ij}^{APC}\right) = -0.5 \times k_B T \ln\left(z_{ij}\right)$$

where T is the temperature (310.15 K) and $k_B$ is the Boltzmann constant. Recalling that each *(i, j)* correlation denotes a correlation between 3-nt windows, the nucleotide pairs *(i, j+2)*, *(i+1, j+1)*, and *(i+2, j)* receive the $\Delta G_{pm}$ bonus. $\Delta G_{pm}$ bonuses are summed for pairs supported by multiple PAIR-MaP correlations. Note that within *RNAstructure*, $\Delta G_{pm}$ bonuses are applied once for edge base pairs and doubly applied to internal base pairs.

***Experimental quality checks.*** RNAs must be modified at sufficient rates in order to reliably measure and prioritize correlations in PAIR-MaP analysis. *PairMapper* thus performs a quality check to confirm that appropriate levels of modification were achieved during the DMS probing experiment. The median co-mutation rate above background is computed for all pairs of windows, $med\left(\, p_{ij}(\boldsymbol{m},\boldsymbol{m}) - p_{ij,bg}(\boldsymbol{m},\boldsymbol{m})\right)$, where $p_{ij}(\boldsymbol{m},\boldsymbol{m})$ and $p_{ij,bg}(\boldsymbol{m},\boldsymbol{m})$ are joint frequencies of the ***(m, m)*** state for the modified and untreated background samples. Datasets with median co-mutation rates below 0.0005, corresponding to 50 observed ***(m, m)*** events at 100,000 sequencing depth, are flagged as unsuitable for PAIR-MaP analysis.

**DMS reactivity normalization**

Raw DMS reactivity rates for each nucleotide ($r_{raw}$) were computed as the difference between the mutation rates of the DMS-modified sample ($M_+$) and the untreated sample ($M_-$):

$$r_{raw} = M_+ - M_-$$

Reactivities of A/C nucleotides and U/G nucleotides were normalized separately by dividing by the average reactivity of the 90th-99th percentile most highly reactive nucleotides:

$$r(n_i) = \begin{cases} r_{raw}(n_i) \;/\; \langle r_{raw}(A,C) \rangle_{[90,99]} \;\; ; \;\; n_i \in (A,C) \\ r_{raw}(n_i) \;/\; \langle r_{raw}(U,G) \rangle_{[90,99]} \;\; ; \;\; n_i \in (U,G) \end{cases}$$

This normalization scheme places A/C and U/G nucleotides on similar 0 to ≈2 reactivity scales and normalizes for experiment-to-experiment variability in overall modification rate. For example, for RMRP, cell-free normalization factors for A/C and U/G nucleotides were 0.148 and 0.018, respectively, and in-cell normalization factors were 0.159 and 0.018, respectively. Normalized DMS reactivities are automatically output by *PairMapper* as text files with the *.dms* suffix.

**Nucleotide-specific DMS reactivity folding restraints**

Normalized DMS reactivities are used to restrain RNA structure modeling by transforming the reactivities into pseudo-energies that favor or disfavor base pairing. Building on a strategy previously used to derive A/C-specific DMS pseudo-energy potentials (24), we obtain nucleotide-specific pseudo-energies ($\Delta G_{DMS}$) from the log-likelihood ratio of a nucleotide being paired versus unpaired given its reactivity:

$$\Delta G_{DMS}(r_i) = -0.5 \times k_B T \ln \left( \frac{P(r_i \mid i \text{ paired}, \; i = X)}{P(r_i \mid i \text{ unpaired}, \; i = X)} \right)$$

T is the temperature (310.15 K), $k_B$ is the Boltzmann constant, $r_i$ is the normalized reactivity of nucleotide *i*, and *X* is a specific nucleotide type (A, C, U, or G). The 0.5 factor accounts for the fact that the $\Delta G_{DMS}$ energy is applied twice for internal base pairs in *RNAstructure* (15). The likelihood functions *P( r | paired, X)* and *P( r | unpaired, X)* were expressed as two-component gamma mixture models fit to reactivity histograms built from our cell-free 23S rRNA dataset.

Reactivity histograms for paired nucleotides were constructed using only doubly stacked pairs (closing base pair nucleotides were excluded). Histograms for unpaired nucleotides were constructed using all single-stranded nucleotides (not Watson-Crick or G-U paired). Fitting was done using the *gammamixEM* function of the *mixtools* package in R. These 23S-rRNA-derived pseudo-energies serve as universal parameters and were used to model folding of all other RNAs, which were not themselves used to develop pseudo-energy terms. We also explored alternative parameterizations, such as including junction base pairs in the paired histograms, excluding the 0.5 prefactor from the $\Delta G_{DMS}$ potential, or using the previously published A/C parameters (24), and found that all parameterization schemes yielded similar structure modeling accuracies (Table S2, and data not shown). Nevertheless, we favor the parameterization described above as the fairest approach for balancing experimental DMS reactivity data and thermodynamic Turner parameters during RNA structure modeling.

**Structure modeling**

*RNAstructure* (v6.0.1) was modified to include the new DMS pseudo-energy potentials described above, which can be accessed using the *-dmsnt* flag in both the *Fold* and *ShapeKnots* programs (15, 25, 26). The *ShapeKnots* code was also updated to support input of single-stranded folding constraints for iterative folding of RNAs with multiple pseudoknots. Both of these upgrades will be made available in future releases of *RNAstructure*.

Structure modeling of the 16S and 23S rRNAs was performed using *Fold* with the *-mfe* and *-md 600* options. For all other molecules, structure modeling was done using *ShapeKnots* with the *-m 1* option. Because *ShapeKnots* is limited to predicting one pseudoknot at a time, *ShapeKnots* modeling was performed iteratively, constraining predicted pseudoknots to be single-stranded (using the *-c* option) and repeating *ShapeKnots* modeling until no further pseudoknots were found. This iterative folding strategy builds on the core strategy used by the *ShapeKnots* algorithm (26). DMS reactivity restraints were passed using the *-dmsnt* option and PAIR-MaP base-pairing restraints were passed using the *-x* option.

**Analyzing sensitivity and positive predictive value**

***PAIR-MaP correlations.*** A principal PAIR-MaP correlation between nucleotide windows [$i$, $i+2$] and [$j$, $j+2$] was considered to be a true positive if [$i\pm1$, $i+2\pm1$] and [$j\pm1$, $j+2\pm1$] were paired in the reference structure (allowing a one-position register shift). In cases where the reference structure contained a two base-pair helix, PAIR-MaP correlations were considered true positives if [$i\pm1$, $i+1\pm1$] and [$j\pm1$, $j+1\pm1$] were paired in the reference. Positive predictive value (ppv) was computed as $ppv = TP/N_{pm}$, where $TP$ is the number of true positives and $N_{pm}$ is the total number of principal PAIR-MaP correlations. PAIR-MaP sensitivity (sens) was computed on a per-helix basis as $sens = h_{pm}/h_{tot}$, where $h_{pm}$ is the number of PAIR-MaP detected helices and $h_{tot}$ is the total number of helices in the reference structure. Reference helices were considered detected if represented by at least one true positive PAIR-MaP correlation. RNA regions lacking PAIR-MaP data (for example, primer binding sites) were excluded from sens and ppv calculations.

***Structure models.*** The sens and ppv of structure models were computed using all WC/GU pairs (including structural regions without data) allowing for one-position register shifts (15).

***Reference structures.*** Accepted reference structures were obtained from refs. (27-31). Two slight revisions were made to the published tmRNA reference structure: (i) consistent with other studies (32), we treat the coding region stem in the region G87-C98 as part of the reference structure; and (ii) our DMS data support pairing of G43-U308 versus G43-U311 typically drawn in the covariation structure (both pairs are equally compatible with covariation data). The RMRP reference structure was revised to include helix P7 identified in our current study (Fig. 5). Pseudoknot-free structures were used for the *E. coli* rRNAs (29). Regions of the 16S and 23S rRNAs with inconsistent DMS reactivity and PAIR-MaP signals were marked as misfolded and excluded from ppv/sens calculations (Fig. 3A, S4). Reference structures for all RNAs, including the 16S and 23S rRNAs with excluded misfolded regions, are available as supporting datasets.

**Covariation analysis**

***RMRP.*** The RNA Families Database (RFAM) provides a structural alignment for RMRP (RF00030) but the alignment is poorly defined in the central junction region (33). We realigned the RFAM sequences against our PAIR-MaP guided human RMRP structure. A Stockholm file containing the human RMRP sequence and in-cell structure model was used to create an initial covariation model (CM) using the *cmbuild* command of Infernal (v1.1.2) (34). The RFAM seed alignment was downloaded as an ungapped fasta and aligned against this initial CM using *cmalign*. Poorly aligning sequences with bitscores < 10 were removed, resulting in a reduced set of 41 ungapped seed sequences (reducedseed.fa). reducedseed.fa was realigned to the initial CM using *cmalign* with *--mapali* and *--mapstr* options, and the resulting alignment was used to build a refined CM. Finally, the complete ungapped RFAM sequence listing (933 sequences) was aligned against the refined CM using *cmalign* with *--mapali* and *--mapstr* options.

***S2-binding motif.*** A structural alignment of the S2-binding motif has been previously published, but only covers the primary stem and pseudoknot (5). We therefore generated a new alignment that includes the full three-way junction. Using a previously described procedure (35), an initial covariation model was built using the *E. coli* structure and trained through iterative Infernal alignments to a bacterial genomic database (database details are described in (35)). Similar to previous studies, we found that the pseudoknot motif is broadly conserved upstream of *rpsB* (5). However, neither the short stem 5' to the primary stem or the closing the helix of the three-way junction was conserved outside of the order *Enterobacteriales*. Analysis of all 32 non-endosymbiont *Enterobacteriales* sequences confirmed that all sequences can form at least a three base pair P1 stem, but no significant covariation is observed (not shown).

***S4-binding motif.*** RFAM provides a multiple sequence alignment for the double pseudoknot structure of the S4-binding motif (RF00140) (33); a more carefully curated alignment was also published by Fu et al. (5). We realigned the Fu sequences against the proposed *E. coli* kissing-loop structure following a similar strategy as used for RMRP. A Stockholm file containing the *E. coli* kissing-loop structure model was used to create an initial CM using the *cmbuild* command of Infernal (34). The Fu alignment was converted to an ungapped fasta and aligned against the initial CM using *cmalign* with *--notrunc --mapali* and *--mapstr* options. Sequences with bitscores < 10 were removed, followed by realignment and building of a refined CM from the reduced set of well-aligning sequences. The complete list of Fu sequences was then aligned against the refined CM using *cmalign* with *--notrunc --mapali* and *--mapstr* options and manually adjusted to ensure proper alignment of PK1 sequences. Due to overlap with the *rpsM* coding sequence, there is minimal sequence variation and hence no covariation signal for either the kissing-loop or the double pseudoknot structure. However, the H3 stem is more highly conserved than the putative double pseudoknot interactions (Fig. 6C, compare to ref. (5)). We also realigned the set of RFAM sequences using a CM built from our complete Fu realignment; this analysis revealed that the H2 stem was also conserved among the larger set of RFAM sequences (not shown).

For all motifs, covariation was assessed using R-scape (v0.7.3) with default parameters (21), and conservation was assessed using R2R with 10% tolerance for non-canonical base pairs, 90% nucleotide conservation threshold, and 50% nucleotide present threshold (36).

### Figure S1

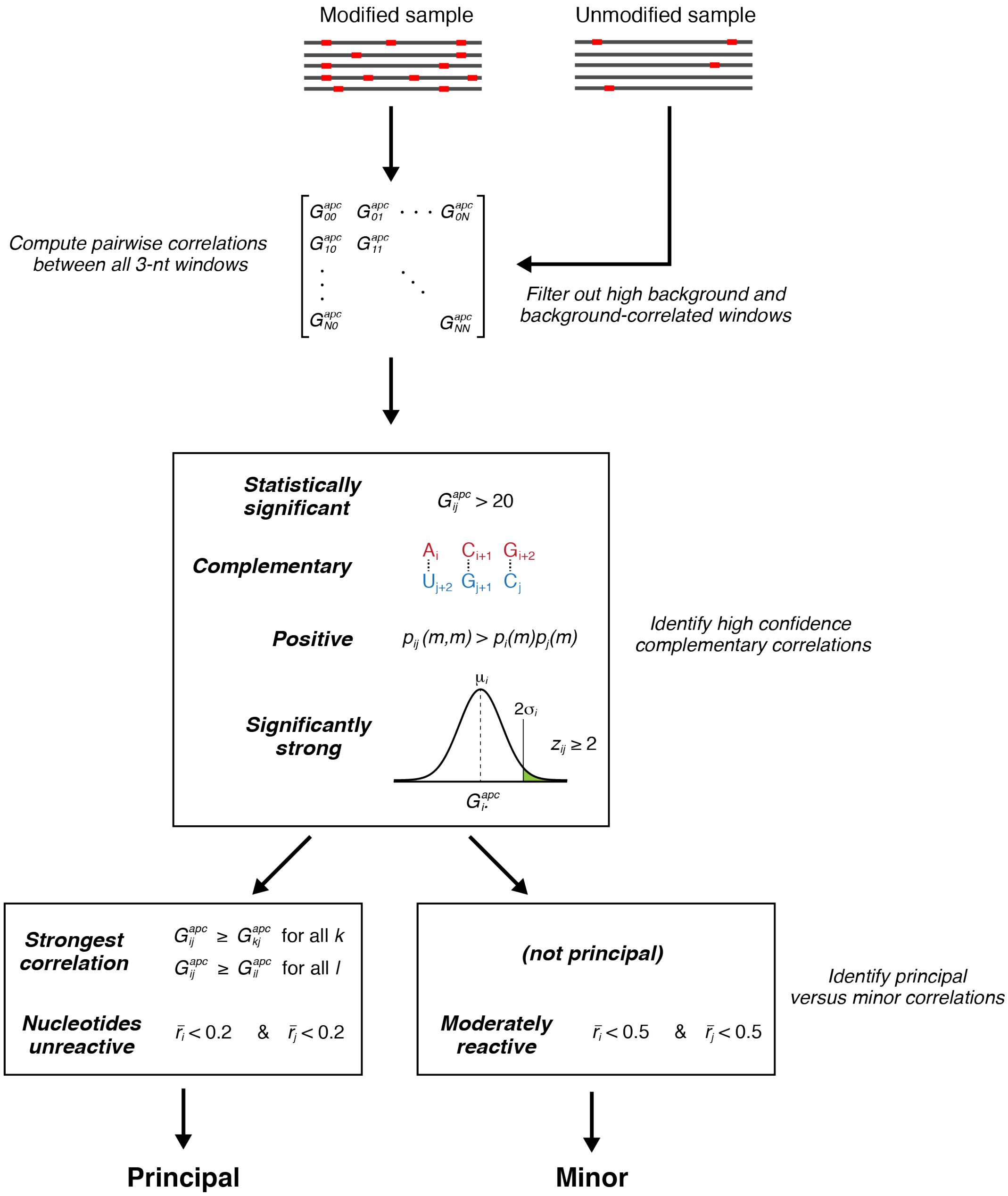


**Figure S1: Outline of the PAIR-MaP strategy.**

**Figure S2**

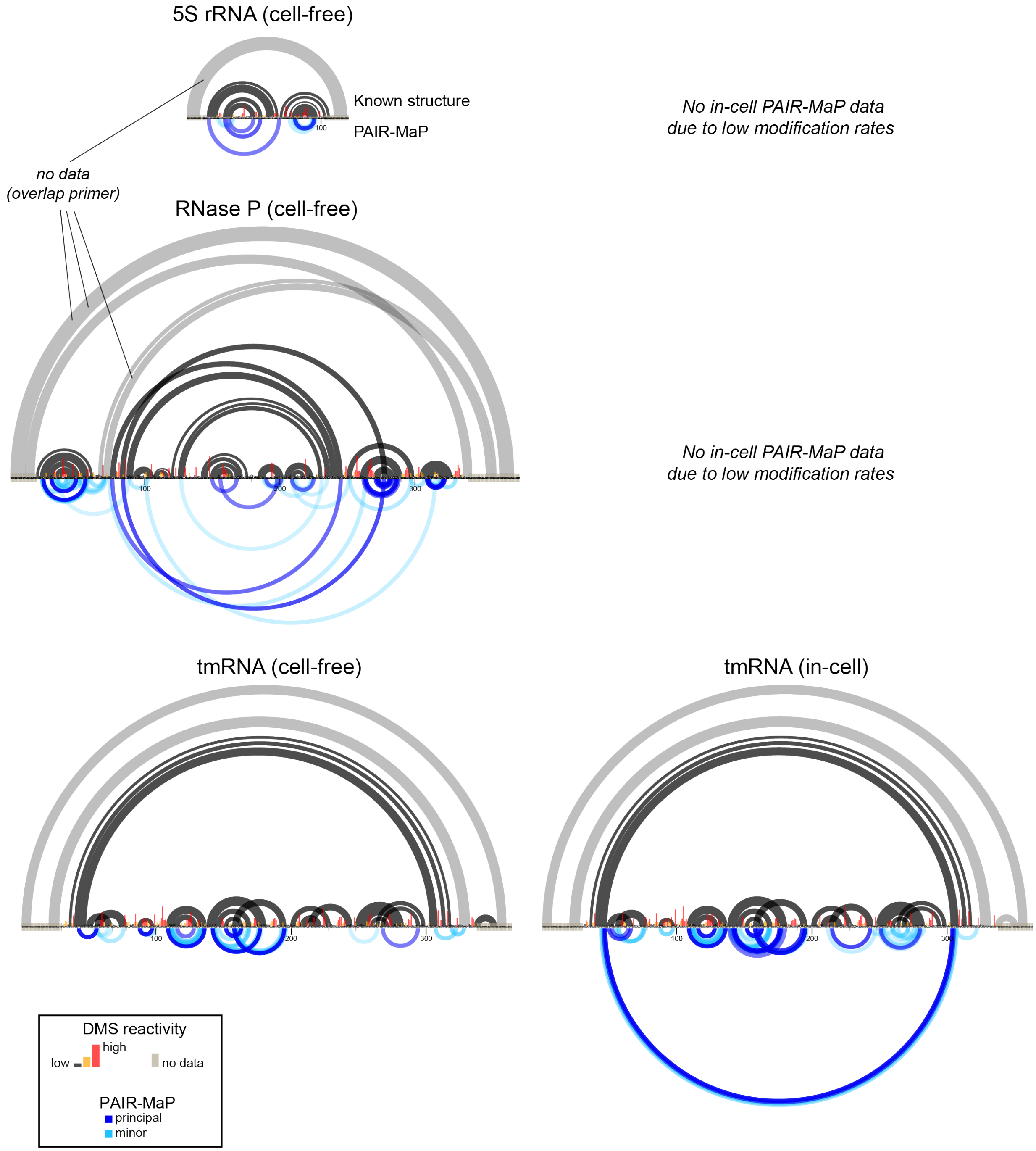


**Figure S2: PAIR-MaP data for *E. coli* small RNAs probed under cell-free and in-cell conditions.** Known secondary structures are shown at top (in gray) and PAIR-MaP correlations are shown at bottom (in dark and light blue). Known helices that are colored light gray overlap primer binding sites and are undetectable by PAIR-MaP. tmRNA in-cell PAIR-MaP data are reproduced from Figure 3 to facilitate comparisons between cell-free and in-cell datasets. Pseudoknots can be identified as helices with overlapping arcs.

**Figure S3**

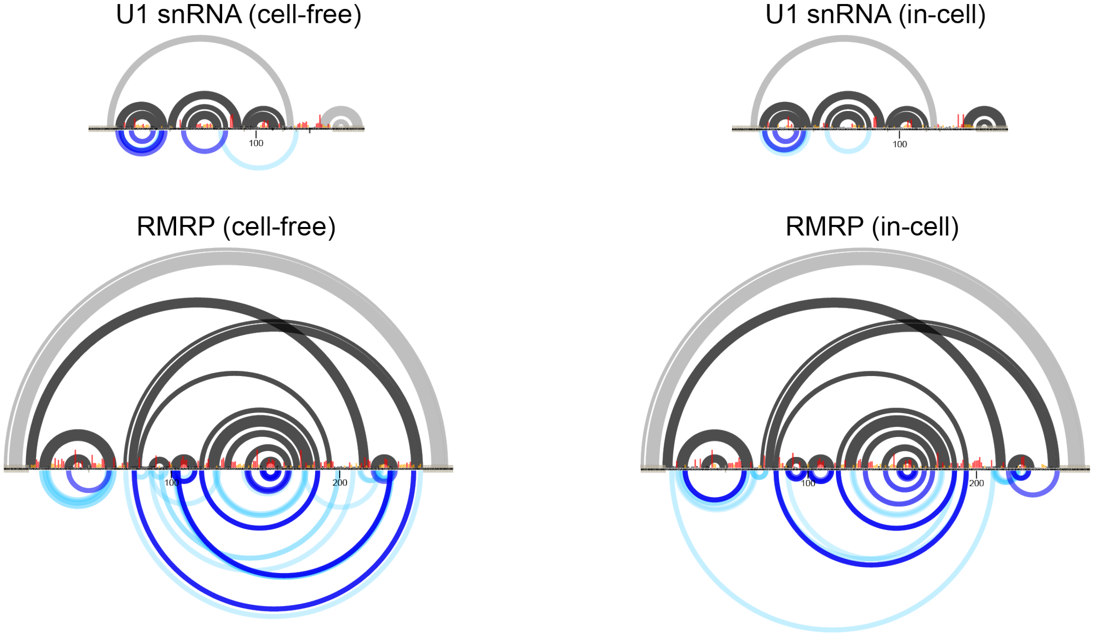


**Figure S3: PAIR-MaP data for human small RNAs probed under cell-free and in-cell conditions.** The key is provided in Figure S2. RMRP in-cell data are reproduced from Figure 5. Misfolding can be identified by non-agreement between PAIR-MaP arcs (bottom) and the accepted structure (top).

### Figure S4

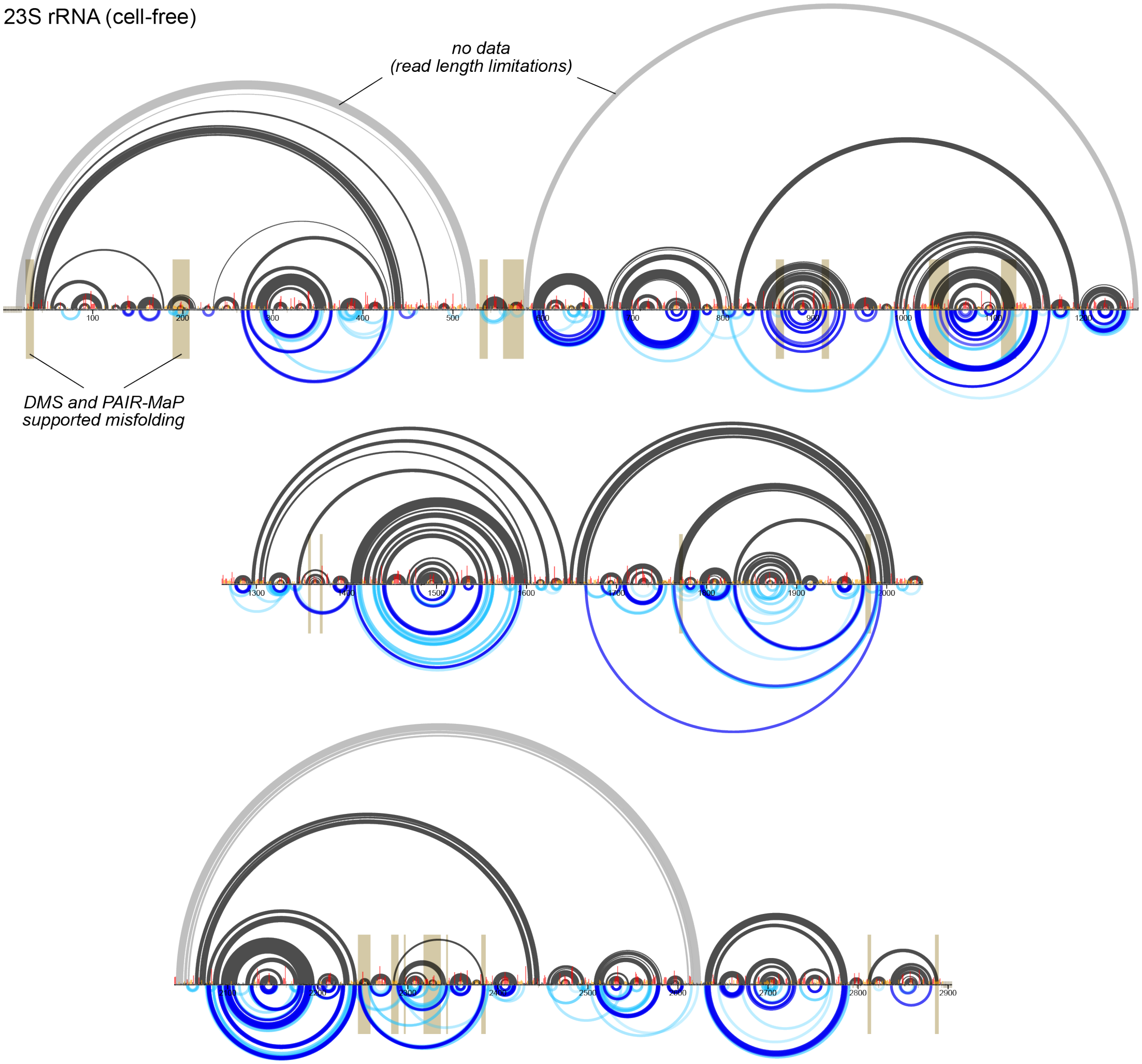


**Figure S4: PAIR-MaP data for *E. coli* 23S rRNA probed under cell-free conditions.** In-cell data were unobtainable due to low DMS modification rates. The three images shown correspond the entire 2904 nt RNA; from top to bottom, domains I and II, III and IV, and V and VI. The long-range Helix 1 is not shown for clarity. The key is provided in Figure S2. Known helices colored light gray span greater than 500 nts and are undetectable by PAIR-MaP due to sequencing read length limitations. Known helix 1 (pairing between nts 1-8 and 2895-2902) is not shown for clarity.

**Figure S5**

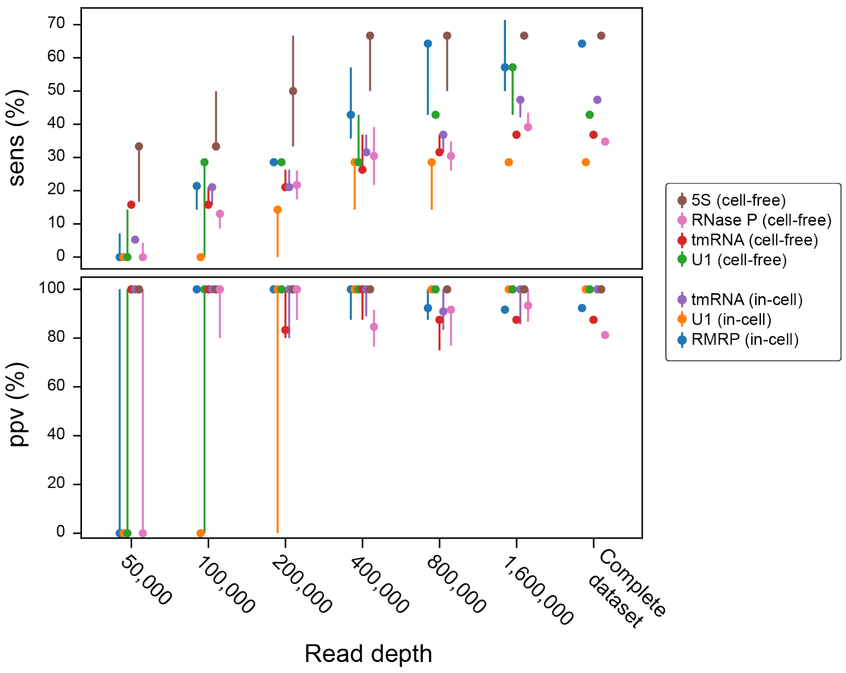


**Figure S5: Dependence of PAIR-MaP accuracy on sequencing depth.** Reads were randomly sampled without replacement to the indicated depth from complete PAIR-MaP sequencing datasets. The ppv and sens of principal PAIR-MaP correlations were computed for five independent samples at each depth, with the median and range shown using dots and whiskers. Sampling was done using the --*undersample* option of *pairmapper.py*. ppv and sens were computed as described in the Methods.

### Figure S6

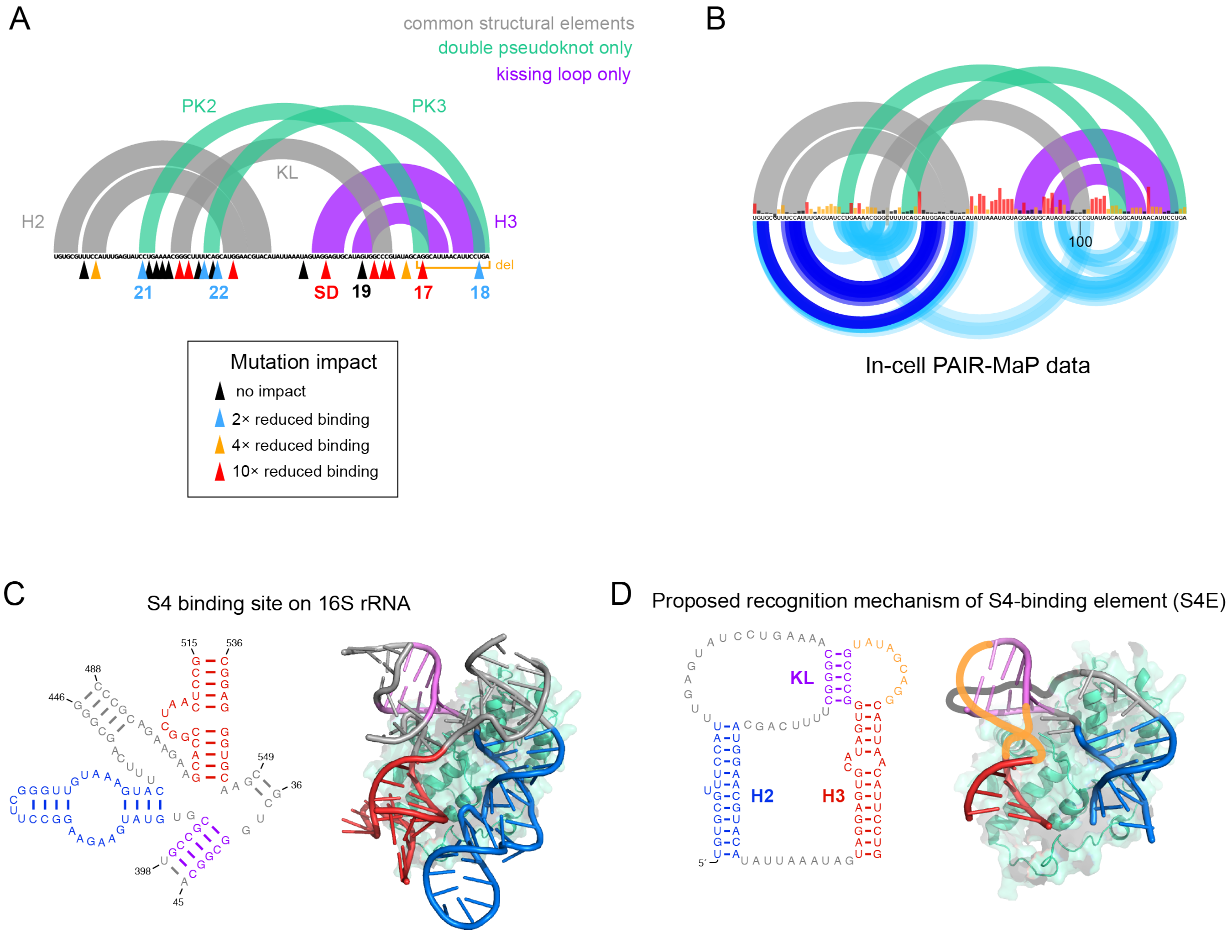


**Figure S6: Proposed S4-binding element (S4E) structures.** (**A**) Map of prior mutational experiments on the S4E. Approximate mutation sites are indicated by arrows and colored by the impact of the independent mutation on S4 binding affinity (1, 4). Mutations discussed in the text are numbered. The previously proposed double pseudoknot structure and the currently proposed kissing loop structure are both shown at top for context. (**B**) Comparison of in-cell PAIR-MaP data with the double pseudoknot and kissing loop structures. (**C**) Secondary structure (left) and crystal structure (right, PDB 4YBB) of S4 binding site on the 16S rRNA. (**D**) Secondary structure diagram of the proposed S4E kissing loop structure (left) and cartoon showing potential homology of the kissing loop structure to the 16S rRNA (right).

**Table S1:** Area under the ROC curve (AUC) values for ROC plots shown in Figure 1D of DMS reactivities. AUC > 0.5 indicates DMS reactivity positively discriminates single-stranded versus base-paired nucleotides, with AUC = 1.0 indicating perfect discriminatory power. AUC = 0.5 indicates DMS reactivity performs no better than random at identifying single-stranded nucleotides.

**Cell-free**

| | Cytosine | Adenosine | Uridine | Guanosine |
|---|---|---|---|---|
| *E. coli* ncRNAs | 0.98 | 0.79 | 0.86 | 0.59 |
| *E. coli* rRNAs | 0.92 | 0.84 | 0.85 | 0.72 |
| Human ncRNAs | 0.91 | 0.87 | 0.85 | 0.60 |
| 1M7 (*E. coli* rRNA) | 0.81 | 0.80 | 0.78 | 0.80 |

**In-cell**

| | Cytosine | Adenosine | Uridine | Guanosine |
|---|---|---|---|---|
| *E. coli* ncRNAs | 0.96 | 0.78 | 0.88 | 0.60 |
| *E. coli* rRNAs | 0.82 | 0.58 | 0.70 | 0.48 |
| Human ncRNAs | 0.94 | 0.79 | 0.84 | 0.55 |

**Table S2:** Comparison of structure modeling accuracy using different DMS pseudo-energy parameterizations. Results from PAIR-MaP+DMS modeling and without experimental data (no data) are reproduced from Table 1. Modeling was also performed using solely the nucleotide-specific DMS parameters described here (Current), and using previously published parameters that only consider A/C nucleotides (Cordero (24)). ppv and sens values for SHAPE-directed rRNA structure models are also provided as a reference (18). Results are colored on a scale to reflect low (red) to high (green) modeling accuracy. RNAs with known misfolding are colored grey and are excluded from averages.

| | PAIR-MaP + DMS | | DMS (Current) | | DMS (Cordero) | | No data | | SHAPE | |
|---|---|---|---|---|---|---|---|---|---|---|
| **Cell-free** | ppv | sens | ppv | sens | ppv | sens | ppv | sens | ppv | sens |
| 16S rRNA | 79 | 85 | 80 | 83 | 80 | 85 | 46 | 51 | 77 | 83 |
| 16S (excl. misfold) | 90 | 96 | 89 | 91 | 92 | 96 | 58 | 66 | 90 | 94 |
| 23S rRNA | 79 | 85 | 81 | 83 | 77 | 83 | 53 | 54 | 76 | 82 |
| 23S (excl. misfold) | 83 | 89 | 84 | 87 | 80 | 87 | 61 | 68 | 79 | 86 |
| 5S rRNA | 82 | 87 | 82 | 87 | 82 | 87 | 25 | 26 | | |
| RNase P | 87 | 87 | 88 | 87 | 86 | 87 | 64 | 67 | | |
| tmRNA | 88 | 90 | 96 | 93 | 91 | 90 | 63 | 58 | | |
| U1 snRNA | 88 | 98 | 88 | 98 | 86 | 98 | 88 | 98 | | |
| RMRP | 74 | 74 | 72 | 72 | 68 | 71 | 58 | 63 | | |
| ***Average*** | 86 | 91 | 88 | 91 | 86 | 91 | 60 | 64 | | |
| **In-cell** | | | | | | | | | | |
| 16S rRNA | n/a | n/a | 75 | 83 | 69 | 78 | 46 | 51 | | |
| 23S rRNA | n/a | n/a | 56 | 65 | 50 | 59 | 58 | 66 | | |
| 5S rRNA | n/a | n/a | 82 | 87 | 82 | 87 | 25 | 26 | | |
| RNase P | n/a | n/a | 85 | 87 | 89 | 93 | 64 | 67 | | |
| tmRNA | 99 | 97 | 94 | 91 | 91 | 90 | 63 | 58 | | |
| U1 snRNA | 91 | 96 | 91 | 96 | 90 | 98 | 88 | 98 | | |
| RMRP | 97 | 97 | 95 | 100 | 95 | 100 | 58 | 63 | | |
| ***Average all*** | | | 83 | 87 | 81 | 86 | 57 | 61 | | |
| ***Average (PAIR-MaP RNAs)*** | 96 | 97 | 93 | 96 | 92 | 96 | 70 | 73 | | |

### Table S3: Oligonucleotides used in this study

| RNA | Sequence (5′ → 3′) |
|---|---|
| Adenine riboswitch | Template:<br>TAATACGACTCACTATAGGCCTTCGGGCCAAGATCAACGCTTCATATAATCCTA<br>ATGATATGGTTTGGGAGTTTCTACCAAGAGCCTTAAACTCTTGATTATGAAGTC<br>TGTCGCTTTATCCGAAATTTTATAAAGAGAAGACTCATGAATTCGATCCGGTTC<br>GCCGGATCCAAATCGGGCTTCGGTCCGGTTC<br><br>PCR forward primer (template amplification):<br>TAATACGACTCACTATAGGCCTTC<br><br>PCR reverse primer (template amplification):<br>GAACCGGACCGAAGCC<br><br>RT primer:<br>Same as PCR reverse primer<br><br>Step1 forward primer:<br>GACTGGAGTTCAGACGTGTGCTCTTCCGATCTNNNNNGGCCTTCGGGCCAA<br><br>Step1 reverse primer:<br>CCCTACACGACGCTCTTCCGATCTNNNNNGAACCGGACCGAAGCC |
| *E. coli* 5S rRNA | RT primer:<br>ATGCCTGGCAGTTCC<br><br>Step1 forward primer:<br>GACTGGAGTTCAGACGTGTGCTCTTCCGATCTNNNNNCTGGCGGCCGTA<br><br>Step1 reverse primer:<br>CCCTACACGACGCTCTTCCGATCTNNNNNATGCCTGGCAGTTCC |
| *E. coli* tmRNA | RT primer:<br>GAGCTGGCGGGAGTTGAA<br><br>Step1 forward primer:<br>CCCTACACGACGCTCTTCCGATCTNNNNNCTGGATTCGACGGGATTTGC<br><br>Step1 reverse primer:<br>GACTGGAGTTCAGACGTGTGCTCTTCCGATCTNNNNNGAGCTGGCGGGAGTTG<br>AA |
| *E. coli* RNase P | RT primer:<br>ATAAGCCGGGTTCTGTCGTG<br><br>Step1 forward primer:<br>CCCTACACGACGCTCTTCCGATCTNNNNNGAAGCTGACCAGACAGTCGC<br><br>Step1 reverse primer:<br>GACTGGAGTTCAGACGTGTGCTCTTCCGATCTNNNNNATAAGCCGGGTTCTGTC<br>GTG |

| | |
|---|---|
| Human U1 snRNA | RT primer:<br>CAGGGGAAAGCGCGAA<br><br>Step1 forward primer:<br>GACTGGAGTTCAGACGTGTGCTCTTCCGATCTNNNNNATACTTACCTGGCAGGG<br><br>Step1 reverse primer:<br>CCTACACGACGCTCTTCCGATCTNNNNNCAGGGGAAAGCGCGAA |
| Human RMRP | RT primer:<br>ACAGCCGCGCTGAGA<br><br>Step1 forward primer:<br>GACTGGAGTTCAGACGTGTGCTCTTCCGATCTNNNNNGTTCGTGCTGAAGGC<br><br>Step1 reverse primer:<br>CCTACACGACGACGCTCTTCCGATCTNNNNNACAGCCGCGCTGAGA |
| *E. coli rpsB* 5′-UTR | RT primer:<br>TGAACACCAGCCTTGAGCAT<br><br>Step1 forward primer:<br>CCCTACACGACGCTCTTCCGATCTNNNNNGGACTTCCGATCCATTTCGT<br><br>Step1 reverse primer:<br>GACTGGAGTTCAGACGTGTGCTCTTCCGATCTNNNNNTGAACACCAGCCTTGAGCAT |
| *E. coli rpsM* 5′-UTR | RT primer:<br>CAGGATGGCTTTAGAACGGGT<br><br>Step1 forward primer:<br>CCCTACACGACGCTCTTCCGATCTNNNNNGCATATTTTTCTTGCAAAGTTGGGT<br><br>Step1 reverse primer:<br>GACTGGAGTTCAGACGTGTGCTCTTCCGATCTNNNNNCAGGATGGCTTTAGAACGGGT |
